# A novel mitragynine analog with low efficacy mu-opioid receptor agonism displays antinociception with attenuated adverse effects

**DOI:** 10.1101/2021.04.22.440994

**Authors:** Soumen Chakraborty, Jeffrey F. DiBerto, Abdelfattah Faouzi, Sarah M. Bernhard, Anna M. Gutridge, Steven Ramsey, Yuchen Zhou, Davide Provasi, Nitin Nuthikattu, Rahul Jilakara, Melissa N.F. Nelson, Wesley B. Asher, Shainnel O. Eans, Lisa L. Wilson, Satyanarayana M Chintala, Marta Filizola, Richard M. van Rijn, Elyssa B. Margolis, Bryan L. Roth, Jay P. McLaughlin, Tao Che, Dalibor Sames, Jonathan A. Javitch, Susruta Majumdar

**Author notes:** Corresponding author(s) : Jonathan A. Javitch, MD PhD, And, Susruta Majumdar, PhD.

## Abstract

Dried kratom leaves are anecdotally used for the treatment of pain, opioid dependence, and alcohol use disorder. We have previously shown that kratom’s natural products (mitragynine) and semi-synthetic analogs (7-hydroxy mitragynine (7OH) and mitragynine pseudoindoxyl) are mu opioid receptor (MOR) agonists that show minimal β-arrestin2 recruitment. To further investigate the structure activity relationships of G-protein potency, efficacy, and β-arrestin2 recruitment, we diversified the mitragynine/7OH templates at the C9, -10 and -12 positions of the aromatic ring of the indole moiety. Three lead C9 analogs, synthesized by swapping the 9-methoxy group with varied substituents, namely phenyl (**SC11**), methyl (**SC12**), 3’-furanyl (**SC13**), were further characterized using a panel of *in vitro* and *ex vivo* electrophysiology assays. All three compounds were partial agonists with lower efficacy than both DAMGO and morphine in heterologous G-protein assays and synaptic physiology. **SC11-13** also showed lower recruitment of both β-arrestin subtypes compared to DAMGO, and in assays with limited MOR receptor reserve, the G-protein efficacy of **SC11, SC12** and **SC13** was comparable to buprenorphine. In mouse models, at equianalgesic doses **SC13** showed MOR-dependent analgesia with potency similar to morphine without respiratory depression, hyperlocomotion, constipation, or place conditioning. Taken together, these results suggest that MOR agonists with a G-protein efficacy profile similar to buprenorphine can be developed into opioids that are effective analgesics with greatly reduced liabilities.

## INTRODUCTION

Opioids targeting the mu opioid receptor (MOR) are used for the treatment of moderate to severe pain.^1^ However, MOR activation is also associated with serious side effects such as tolerance, physical dependence, and risk of abuse;^1–3^ opioid-induced respiratory depression can be lethal at high doses and constipation can be debilitating as well. Opioid abuse and overdose are one of the leading causes of accidental death in the United States, responsible for more than 47,000 deaths in 2019 alone.^4^ Therefore, discovery of a new class of MOR agonist molecular scaffold that retains potent analgesic actions but with reduced side effects and abuse potential is a pressing challenge for the scientific community.

Applying molecular modeling based on active state MOR structures, synthesis of novel ligands and using newer assays with limited receptor reserve, the opioid field is revisiting the strategy of developing low efficacy partial agonists as novel safer analgesics.^5–7^ Numerous MOR partial agonists with multifunctional activity at other opioid receptor subtypes have been described in the literature, such as buprenorphine, nalbuphine and pentazocine, validating the feasibility of this strategy. The identification of novel partial agonists may have been hindered by modern screening assays that assess G-protein activity, yet have large receptor reserve (so called “spare” receptors), which prevents a simple delineation of lower efficacy compounds.^8, 9^ In order to develop candidate pain relievers based on mitragynine and 7OH scaffolds with a particular goal of assessing G-protein efficacy and its impact on opioid function *in vivo*, we aimed at diversification of the mitragynine template and evaluated the resulting compounds in systems capable of detecting their true efficacy.

The psychoactive plant *Mitragyna speciosa*, commonly known as kratom, has traditionally been used for the treatment of opioid dependence.^10^ The dry leaves of kratom are used in traditional medicine as an analgesic treatment and are typically consumed directly or brewed as tea. The major active alkaloid found in kratom is mitragynine, along with more than 40 other minor alkaloids.^11–16^ In recent years, we have become interested in the chemistry and pharmacology of kratom alkaloids as probes to understand opioid receptor function.^11, 17–23^ Previous reports from our group reported that mitragynine (possessing an indole core), its oxidation product 7OH (possessing an indolenine core), and mitragynine pseudoindoxyl (**MP**, a skeletal rearrangement product of 7OH with a spiro-pseudoindoxyl core) (**Figure 1A**), are all opioid antinociceptive agents^18, 19^ and G-protein biased MOR agonists.^17, 18, 20, 22^ We also reported oxidative metabolism of mitragynine to 7OH mitragynine using a CYP3A-mediated pathway following oral administration of mitragynine in mice.^19^ Metabolism of mitragynine to 7OH *in vitro*^24^ and in dogs^25^ has been reported by other groups too. More recently, we also reported an atomic-level description of how kratom alkaloids may bind and allosterically modulate MOR^21^. *In vivo* studies in mice revealed that kratom and a number of its alkaloids are analgesic^16, 18, 19, 26–28^ ameliorate opioid physical dependence,^23, 26^ decrease alcohol intake^22^, and inhibit self-administration of heroin in rats.^29^ While 7OH retains its abuse potential after both intravenous and parental administration^22^, intravenous mitragynine is not self-administered,^29, 30^ suggesting that it may be possible to design a safer analgesic based on this template by further optimization of the mitragynine template.

**Figure 1.**
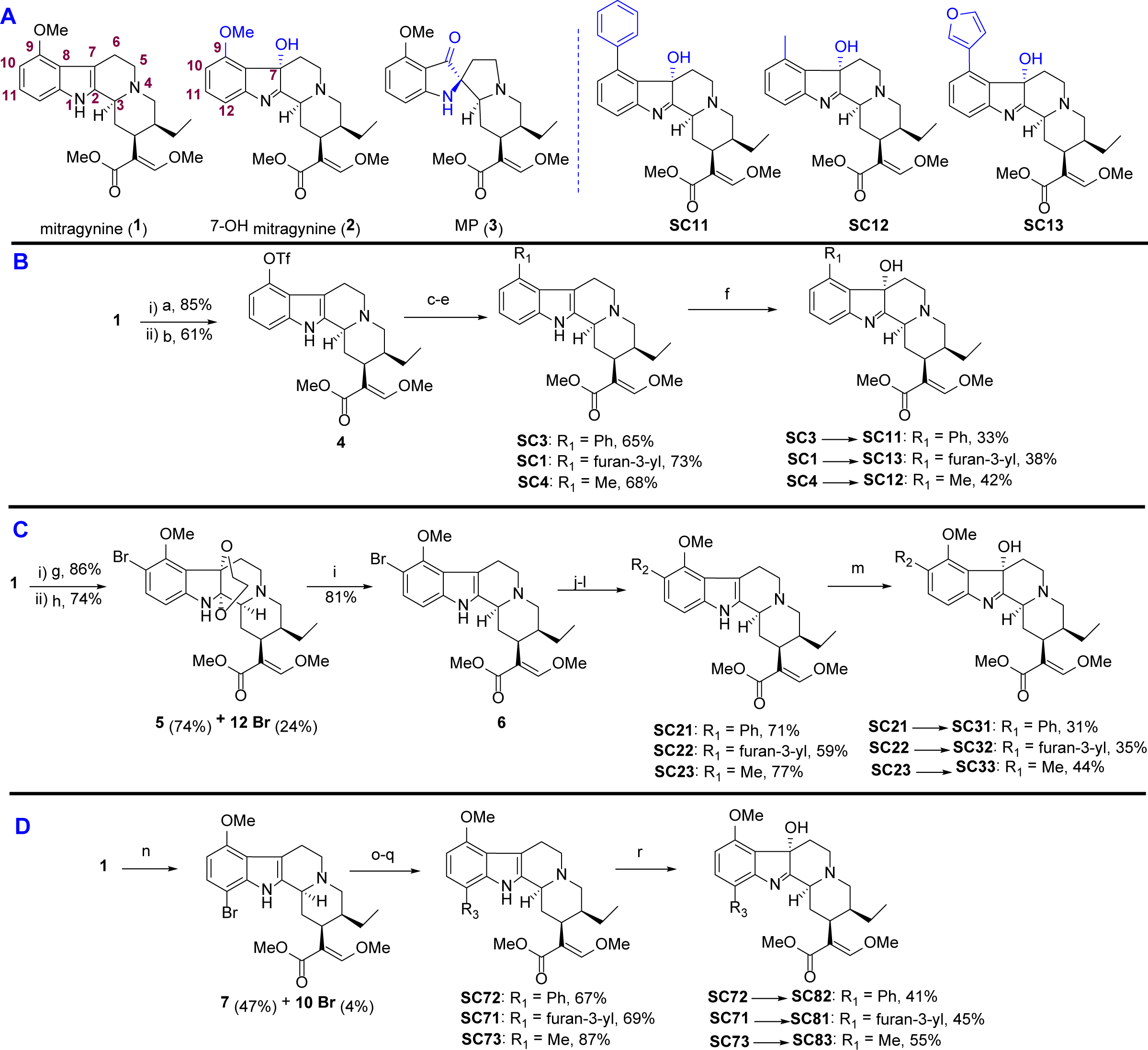
Chemistry. **A) Structure of selected natural and synthetic analogs. B) Synthesis of C9 mitragynine and 7OH derivatives. C) Synthesis of C10 mitragynine and 7OH derivatives. D) Synthesis of C12 mitragynine and 7OH derivatives.** Reagents and conditions: (a) AlCl_3_, EtSH, DCM, 0 °C, 5h ; (b) PhNTf_2_, Et_3_N, DCM, rt, 12h; (c, yielding **SC3**) phenylboronic acid, Pd(PPh_3_)_4_, K_2_CO_3_, MeOH, toluene, 80 °C, 8h; (d, yielding **SC1**) 3-furanylboronic acid, Pd(PPh_3_)_4_, K_2_CO_3_, MeOH, toluene, 80 °C, 8h; (e, yielding **SC4**) DABAL-Me_3_, Pd_2_(dba)_3_, XPhos, THF, 60 °C, 8h; (f, yielding **SC11**, **SC13** and **SC12**) oxone®, NaHCO_3_, H_2_O, acetone, 0 °C, 1h. (g) ethylene glycol, PIFA, CH_3_CN, 0 °C, 1h; (h) NBS, DMF, 5h, rt; (i) NaBH_3_CN, AcOH, MeOH, reflux, 12h; (j, yielding **SC21**) phenylboronic acid, Pd(dppf)Cl_2_, KOAc, THF, 70 °C, 6h; (k, yielding **SC22**) 3- furanylboronic acid, Pd(dppf)Cl_2_, KOAc, THF, 70 °C, 6h; (l, yielding **SC23**) DABAL-Me_3_, Pd_2_(dba)_3_, XPhos, THF, 60 °C, 8h; (m, yielding **SC31**, **SC32** and **SC33**) oxone®, NaHCO_3_, H_2_O, acetone, 0 °C, 1h. (n) NBS, AcOH, 4h, rt; (o, yielding **SC72**) phenylboronic acid, Pd(PPh_3_)_4_, K_2_CO_3_, MeOH, toluene, 80 °C, 8h; (p, yielding **SC71**) 3-furanylboronic acid, Pd(PPh_3_)_4_, K_2_CO_3_, MeOH, toluene, 80 °C, 8h; (q, yielding **SC73**) DABAL-Me_3_, Pd_2_(dba)_3_, XPhos, THF, 60 °C, 8h; (r, yielding **SC82**, **SC81** and **SC83**) oxone®, NaHCO_3_, H_2_O, acetone, 0 °C, 1h.

Chemistry studies to date are limited in the structure-activity relationship (SAR) investigations of both mitragynine and 7OH scaffolds, prompting the present development of diversification strategies across these two indole-based templates. Here, we report the pharmacological characterization of mitragynine and 7OH mitragynine analogs synthesized by introducing a phenyl, 3’-furanyl, and methyl group at the C9/10/12 positions of the aromatic ring in the two templates. The lead compounds **SC11**, **SC12** and **SC13** (**Figure 1A**) showed lower G-protein efficacy at MOR than DAMGO and morphine in *in vitro* assays with limited receptor reserve and *ex vivo* assays as well. The most potent and selective Gi-1 MOR agonist among the three leads, **SC13**, displayed antinociceptive activity comparable to morphine but exhibited greatly attenuated constipation, respiratory depression, and locomotor activity. Furthermore, **SC13** displayed no reinforcement behavior in a conditioned-place preference assay. Taken together, the reported *in vitro* assays in cells*, ex vivo* assays in rat brain slices, and *in vivo* experiments in mouse suggest that the partial agonist **SC13** exerts effective MOR-mediated analgesia with a side effect profile far superior to clinically used MOR-based antinociceptive agents.

## RESULTS

### Chemistry

To assess the pharmacological profile of mitragynine and 7OH templates, structure activity relationships (SAR) studies were carried out by modifying three different regions of the aromatic indole ring on both scaffolds, namely the C9, C10, and C12 positions, with phenyl, 3’furanyl and methyl group substitutions. (**Figure 1**). The unsaturated acrylate segment of both templates is thought to be an essential component for the efficient binding of any mitragynine- or 7OH-related analog into the orthosteric MOR binding pocket.^17^ Therefore, this feature of both scaffolds was kept constant throughout our studies. We synthesized a total of 18 analogs and investigated their pharmacology with *in vitro* assays.

Synthesis of analogs was initiated from mitragynine (**1**) extracted from dry kratom powder following a modified protocol reported by Varadi et al.^18^ To gain access to the C9 position on the mitragynine scaffold, **1** was converted to triflate (**4**, **Figure 1B**). This intermediate triflate was converted to **SC1**, **SC3,** and **SC4** using palladium-catalyzed coupling reactions. Then **SC1**, **SC3,** and **SC4** were transformed to their corresponding 7OH derivatives **SC13**, **SC11,** and **SC12,** respectively, using oxone® and aqueous NaHCO3. Functionalization of the C10 position on the mitragynine scaffold was achieved by selectively incorporating bromine at the C10 position using a protocol developed by Takayama.^31^ Mitragynine (**1**) was first converted to 10-bromo mitragynine **6** using a 3-step reaction sequence (**Figure 1C**). 10-bromo mitragynine **6** was then subjected to different coupling reactions to obtain C10 mitragynine analogs, namely **SC21**, **SC22,** and **SC23**. **SC21-23** were then treated with oxone® and aqueous NaHCO_3_ to obtain the corresponding 7OH derivatives **SC31-33,** respectively. For the C12 derivatives, as shown in **Figure 1D**, mitragynine (**1**) was brominated directly to afford mainly 12-bromo mitragynine (**7**). The same reaction sequence (as in C10) was followed to synthesize C12 substituted analogs **SC71**-**73**. Next, all were treated with oxone® and aqueous NaHCO_3_ to yield C12 7OH derivatives **SC81-83**. Detailed synthetic procedures and characterization data for each of the compounds are shown in the methods section.

### SAR and *in vitro* functional screening of synthesized analogs

Each synthesized compound was first evaluated for G-protein activity using the high throughput Glo-sensor cAMP inhibition assay and Tango β-arrestin2 recruitment assay. For cAMP assays, HEK-T cell lines transiently expressing human MOR, KOR and DOR were used, while for Tango assays, HTLA cells transiently expressing TEV fused β-arrestin2 were used. The data were normalized to that of prototypical full agonists, DAMGO for MOR, U50,488H for KOR, and DPDPE for DOR, respectively. cAMP and β-arrestin2 data for MOR are presented in **Table 1** with representative SAR of selected compounds shown in **Appendix 1-Figure 1A-B**. Additionally, results for KOR and DOR are summarized in **Appendix 1-Table 1** and **Appendix 1-Table 2**.

**Table 1.**
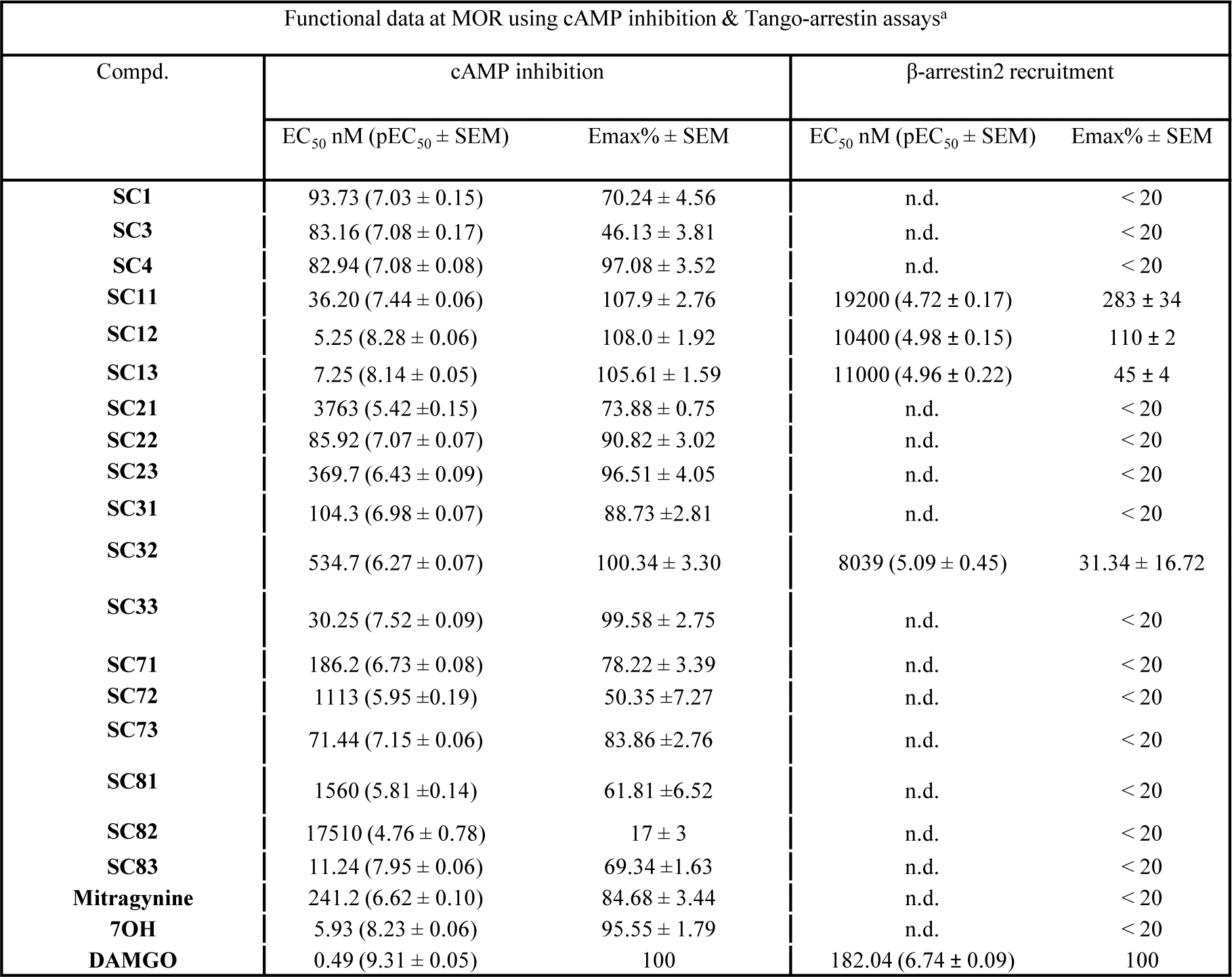
Functional studies at MOR using cAMP inhibition & Tango-arrestin assays. ^a^The functional data of each compound in cAMP and Tango β-arrestin2 in human mu-opioid receptor (hMOR) were determined and normalized to E_max_ of the corresponding standard DAMGO. Results were analyzed using a three-parameter logistic equation in GraphPad Prism and the data are presented as mean EC_50_(pEC_50_ ± SEM) with E_max_% ± SEM for assays run in triplicate. nd; results could not be determined because efficacy of β-arrestin2 recruitment was less than 20%.

We initiated our investigations with modification at the C9 position of mitragynine with the syntheses of **SC1** (9-3’-furanyl mitragynine), **SC3** (9-phenyl mitragynine), and **SC4** (9-methyl mitragynine), each of which revealed moderate activity and potency (EC_50_ > 50 nM) in cAMP assays and poor β-arrestin2 recruitment (E_max_ < 20%) at MOR. We then investigated three other C9 analogs on the 7OH template, 9-3’-furanyl 7OH (**SC13**), 9-phenyl-7OH (**SC11**) and 9-methyl- 7OH (**SC12**). We specifically picked these moieties in order to explore the effect of an aryl- (phenyl), a heteroaryl-(3’-furanyl) and an aliphatic group such as methyl on this template. Incorporation of a phenyl ring at the C9 end of the 7OH scaffold led to an increased cAMP potency at MOR (EC_50_ = 36.2 nM, **Table 1** and **Appendix 1-Figure 1A)** in comparison with the same substituent on the mitragynine template (EC_50_ = 83.2 nM, **SC3**, **Table 1**). Introduction of an aliphatic methyl group at C9 of the 7OH scaffold in **SC12** improved potency in the cAMP assay with MOR (EC_50_ = 5.3 nM) compared to the 9-methyl mitragynine **SC4** (EC_50_ = 82.9 nM). Furthermore, grafting of a 3’-furanyl group at C9 of 7OH (**SC13)** showed similar potency (EC_50_ = 7.3 nM) in the cAMP assay to that of **SC12** as well as the parent 7OH mitragynine (EC_50_= 5.9 nM). Interestingly, while the corresponding analogs on the mitragynine template (**SC1, 3** & **4**) showed poor β-arrestin2 recruitment, the analogs on the 7OH template (**SC11** and **12**) showed robust arrestin recruitment in Tango assays at MOR relative to DAMGO: **SC13** showed 45% β- arrestin2 efficacy relative to DAMGO, but higher than the parent template 7OH (**Table 1** and **Appendix 1-Figure 1B**). The potencies of **SC11, SC12 and SC13** for recruiting arrestin however remained poor (with EC_50_>10 µM for each). **SC11-13** showed no β-arrestin2 recruitment at KOR, but β-arrestin2 recruitment was seen at DOR (E_max_>100%) with all three analogs (**Appendix 1-Table 1** and **Appendix 1-Table 2**). In cAMP assays, **SC13** was most selective for MOR over DOR and KOR, showing 30-fold and 14-fold selectivity compared to **SC12** and **SC11**, which were less MOR selective. (**Table 1**, **Appendix 1-Table 1, and Appendix 1-Table 2**).

The next set of analogs were designed at the C10 and C12 ends of both the mitragynine and 7OH templates. None of the synthesized analogs at C10 and C12 exhibited promising activities at MOR in the cAMP assay except for 12-methyl 7OH (**SC83**), with an EC_50_ = 11.2 nM. Notably, these analogs also did not effectively recruit β-arrestin2 (E_max_ < 20%) in the Tango assay (**Table 1**).

Our mitragynine template diversification attempts did produce numerous partial agonists, but with the exception of **SC83**, their potency was greater than 50 nM, in the cAMP assay. Therefore, **SC11, SC12** and **SC13** (all C9 substituted 7OH analogs) were chosen as leads from the series of compounds synthesized. **SC11-13** were evaluated in the PathHunter assay,^18, 22^ which we and others^12^ have previously used to measure β-arrestin2 activity of the parent natural products. In this assay, like morphine (E_max_=31%), **SC11-13** were found to recruit β-arrestin2 with greatly reduced efficacy (E_max_<20%) compared to DAMGO (**Appendix 1-Figure 1C**). These observations suggest that the much higher β-arrestin2 recruitment seen in the Tango assay is likely a consequence of higher amplification of arrestin signaling compared to the PathHunter assay. In hMOR (human MOR) competition binding assays using ^3^H-DAMGO as the radioligand, DAMGO and morphine showed subnanomolar affinity for MOR; among the lead analogs **SC13** had the highest affinity (K_i_ = 6 nM) and **SC11** and **SC12** had high (15-17 nM) affinity at MOR as well (**Appendix 1-Figure 1D**).

### SC11-13 are MOR partial agonists in BRET-based G-protein activation assays

We next assessed **SC11-13** and the controls DAMGO, morphine, buprenorphine, and fentanyl for G-protein activation (Gi-1) using TRUPATH assays and arrestin recruitment (β-arrestin1/2) using another BRET-based assay, which produce less signal amplification compared to the cAMP and Tango assays.^32^

**SC11-13** showed MOR partial agonist activity with E_max_=60-70% of DAMGO at Gi-1. Fentanyl showed higher efficacy (E_max_=122%) and morphine showed an efficacy only slightly lower than DAMGO (E_max_=94%), whereas buprenorphine had an E_max_=44% in this assay (**Figure 2A** and **Appendix 1-Table 3**). Thus, the intrinsic efficacy of **SC11-13** appeared somewhat higher than for the prototypic MOR partial agonist buprenorphine, but lower than DAMGO, fentanyl and morphine under these conditions (**Appendix 1-Figure 1E** and **Appendix 1-Table 3**).

**Figure 2.**
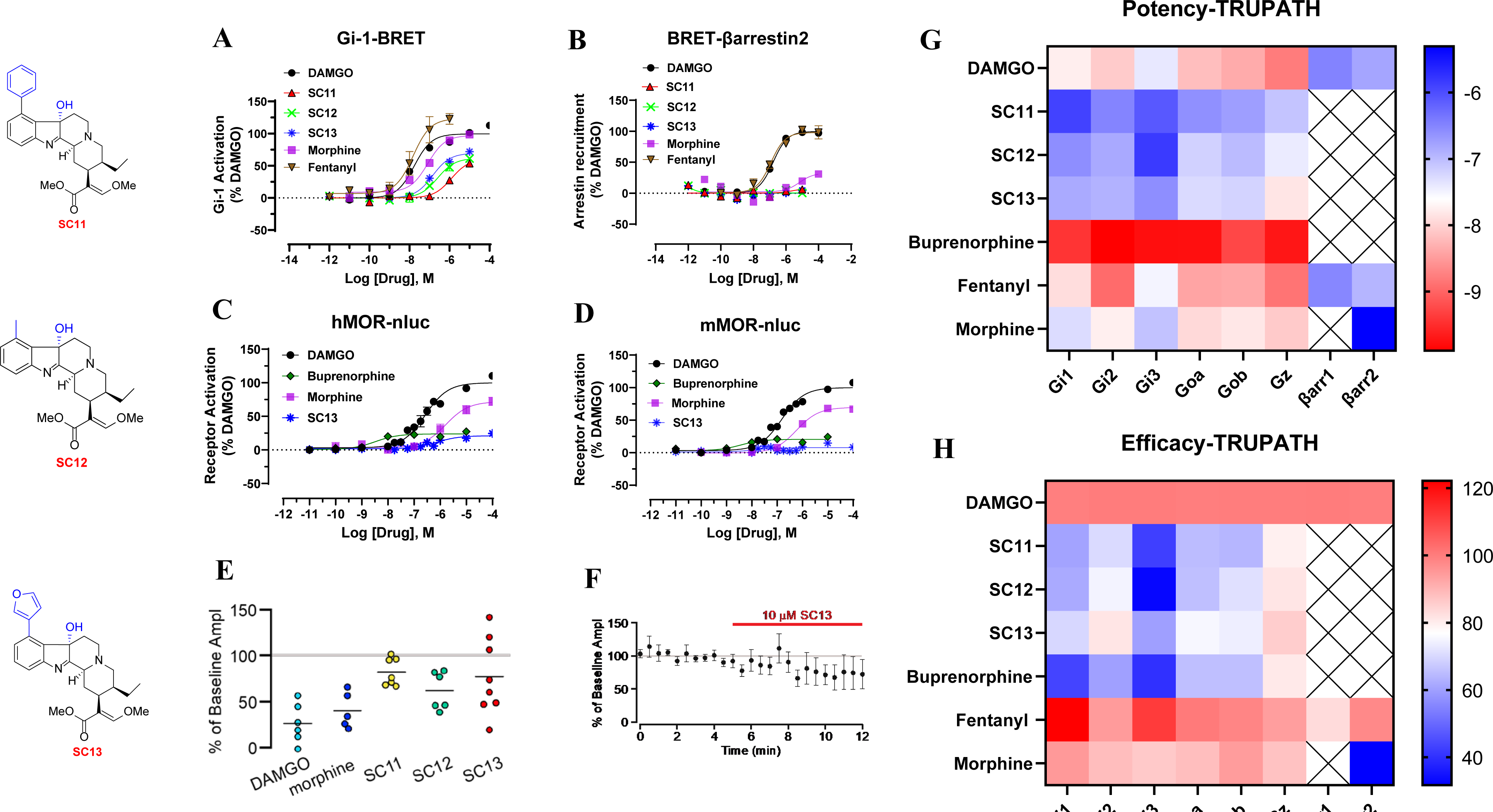
G-protein signaling, arrestin signaling, whole cell electrophysiology in rat VTA and Gα-subtype screening of SC 11-13 and MOR controls in hMOR. SC11-13 are MOR partial agonists in cell based assays G-protein signaling assays as well as in ephys assays. **A) SC 11**-**13** are MOR partial agonists with lower efficacy than morphine, fentanyl and DAMGO in Gi-1 BRET assays. **B) SC11**-**13** showed no measurable β-arrestin2 recruitment (<10%) in BRET arrestin recruitment assays compared to fentanyl and DAMGO in this assay. **C)** In Nb33 recruitment assays measured using BRET assays in hMOR, **SC13** had lower efficacy than DAMGO and morphine and similar efficacy to buprenorphine. **D)** In Nb33 recruitment assays measured using BRET assays in mMOR, **SC13** had lower efficacy than DAMGO and morphine and similar efficacy to buprenorphine. **E)** Summary inhibition of electrically evoked IPSCs in VTA neurons in response to 5 μM DAMGO, 10 μM morphine, 10 μM **SC11**, 10 μM **SC12**, 10 μM **SC13**, where each circle is one neuron. Horizontal bars indicate means. **SC 11-13** show lower efficacy than DAMGO. **F)** Mean time course of responses during bath application of **SC13**, n = 8 in whole cell electrophysiology in rat VTA. See **Appendix 1- Table 3** in **SI** for values for panels A-D. **G)** TRUPATH heatmaps demonstrate how a panel of kratom analogs, **SC11-13** and MOR agonists engage Gαi/o-class transducers with varying potency (**G**) and efficacy (**H**). Most ligands exhibit enhanced (GαZ) relative to other G-protein transducers. Heatmap colors represent mean log(EC50) and normalized efficacy values; NR, no response, presented as a white square. Mean values and standard error are reported in **Appendix 1-Table 4**. Data for all functional assays were carried out in hMOR was normalized to E_max_ of DAMGO. The dose response curves were fit using a three-parameter logistic equation in GraphPad Prism and the data are presented as mean EC_50_(pEC_50_ ± SEM) for assays run in triplicate.

The novel compounds and MOR controls were also characterized using the TRUPATH assay^32^ for activation of other Gα-i/o subtypes (Gi-2, Gi-3, GoA, GoB and Gz). **SC11, SC 12,** and **SC13** were found to be partial agonists at all these G-protein subtypes and showed an efficacy profile similar to buprenorphine at the same subtypes (**Figure 2H** and **Appendix 1-Table 4**). The highest potencies (**Figure 2G** and **Appendix 1-Table 4**) and efficacy (**Figure 2H** and **Appendix 1-Table 4**) were seen at Gz for **SC11-13** as well as the MOR reference compounds. Specifically, the Gz efficacy for **SC11-13** was similar to both buprenorphine and morphine, but lower than DAMGO and fentanyl. Notably, the higher efficacy and equipotency at Gi-1 and Gz for buprenorphine relative to DAMGO and higher potency of morphine at Gz relative to Gi-1, are consistent with recent work from the Bidlack group.^33^

In β-arrestin2 recruitment assays, DAMGO (E_max_=100%) and fentanyl (E_max_=98%) robustly recruited β-arrestin2 (**Figure 2B** and **Appendix 1-Table 3**), whereas morphine was moderately active (E_max_=32%). and buprenorphine was less active (E_max_<10%) (**Appendix 1-Figure 1F** and **Appendix 1-Table 3**). β-arrestin2 recruitment induced by incubation with buprenorphine and **SC11-13** was not measurable with this assay (**Appendix 1-Figure 1F** and **Appendix 1-Table 3**). In this assay, **SC11, SC12, SC13**, morphine, and buprenorphine failed to show recruitment of β-arrestin1, whereas fentanyl displayed 83% efficacy in this assay compared to DAMGO. In summary, in the BRET-based assays, **SC11-13** acted as MOR partial agonists for G protein activation but not show arrestin recruitment.

We also evaluated MOR selectivity of **SC13** (our behavioral lead-see next section) versus KOR/DOR selectivity for Gi-1 activation. **SC13** was found to have ∼100-fold lower potency at DOR and KOR (**Appendix 1-Figure 1G-H)** in this assay.

Recent work have shown that nanobodies (Nb33 and Nb39) can be used as receptor-activation sensors to accurately probe agonist activity.^34, 35^ Canals and co-workers have recently used a conformationally selective Nb33-recruitment assay^20^ to more accurately determine efficacy^5^ of MOR ligands in an unamplified manner more akin to the BRET-based direct arrestin recruitment assay. Since morphine was a partial agonist in this assay (E_max_=71%), compared to 94% in our Gi-1 assays, we used this assay to determine the efficacy of **SC11-13** and compared it to DAMGO, morphine and buprenorphine in HEK293-T cells transiently transfected with the human and murine-MOR. In this assay, the efficacies of morphine and buprenorphine were 72% and 24% respectively compared to DAMGO, and **SC11, 12** and **13** each showed efficacies of ∼20% in this assay at hMOR (**Figure 2C/ Appendix 1-Figure 1I** and **Appendix 1-Table 3**). Similarly at murine MOR (**Figure 2D/ Appendix 1-Figure 1J** and **Appendix 1-Table 3)** the efficacies of **SC11-12** (E_max_=15-18%) and **SC13** (8%) were more comparable to buprenorphine (20%) and lower than morphine (69%). Thus, the efficacies of our lead ligands are similar to buprenorphine and far lower than morphine in this assay, and the efficacies of our control drugs matched published reports.^5^ While it is difficult to accurately determine potencies of our leads with such limited dynamic range, the potency of **SC13** (our behavioral lead) was in the same range as morphine as well as DAMGO **(Figure 2C/D)**.

### SC11-13 showed low efficacies for inhibition of synaptic transmission

To gauge partial agonism in a physiologically natural, endogenous system, we utilized whole cell electrophysiological recordings from ventral tegmental area (VTA) neurons in acute rat brain slices. Full MOR agonists such as DAMGO strongly inhibit GABA release onto VTA neurons (**Figure 2E**).^36^ Thus, we tested the efficacy of 10 μM **SC11, SC12**, and **SC13** at this synapse by measuring electrically-evoked GABA_A_ receptor-mediated IPSCs. As a control, in brain slices from the same rats, we also measured responses to a saturating concentration of DAMGO (5 μM), and 10 µM morphine. The mean inhibition of evoked IPSCs was smaller in response to **SC11, SC12**, and **SC13** compared to DAMGO as well as morphine (**Figure 2E**). The mean time course of the response to 10 µM **SC13** is shown in **Figure 2F.** Together these effects are consistent with the **SC11-13** compounds acting as partial agonists, at this synapse.

### SC11-13 form different interactions with MOR compared to morphine and buprenorphine

To provide a structural context to the observed pharmacological differences between kratom alkaloids and classical opioid drugs, such as morphine and buprenorphine, we carried out a statistical analysis of the interactions formed between MOR residues and each of the compounds included in this manuscript, during molecular dynamics (MD) simulations of ligand-receptor complexes embedded in hydrated 1-palmitoyl-2-oleoyl phosphatidyl choline (POPC) bilayers. DAMGO and fentanyl were excluded from this analysis because of their different chemical composition and expected unique mode of binding with respect to the other molecules in the data set. The MD simulations of **SC1, SC3, SC4, SC11, SC12, SC13, 11F**,^37, 38^ morphine, and buprenorphine (See **Appendix 1-Table 5** for efficacy data), were carried out using the same simulation parameters and protocol used in our previous work on 7OH and mitragynine.^39^ A statistical analysis of structural interaction fingerprints (SIFts) derived from these simulations and whose average probabilities are listed in **Appendix 1-Table 6** for each ligand, yielded 8 statistical models that best recapitulate the negative logarithm of experimental G protein E_max_ values obtained for each ligand (see **Appendix 1-Figure 4**, Gi-1 E_max_ was used for ligands). These models correspond to the top quartile of R^2^ validation on the full training set and the lowest root-mean-square error (RMSE) on the leave-one-out (LOO) validation (red dots in **Appendix 1-Figure 3**). According to this modeling and the calculated average positive coefficients reported in **Appendix 1-Table 7**, the efficacy of **SC11-13** ligands and buprenorphine is reduced because of specific apolar interactions these ligands form with Y75(1.39), N127(2.63), I144(3.29), C217(45.50), and W133(23.50). On the other hand, the efficacy of **SC11-13** is enhanced (negative coefficients) by specific apolar and edge-to-face aromatic interactions these ligands form with H319 (7.36). The aforementioned residue numbers refer to the murine MOR sequence and dot-separated numbers in parenthesis refer to the Ballesteros-Weinstein generic numbering scheme^40^ when located in transmembrane (TM) helices and the Isberg’s numbering scheme^41^ when in loops. The first number in these schemes refers either to a helix (e.g., 3 refers to TM3) or a loop (e.g., “45” refers to the loop between TM4 and TM5) to which that residue belongs, whereas the second number represents the residue position relative to the most conserved residue in the helix, which is always defined by the number 50.

**Figure 3.**
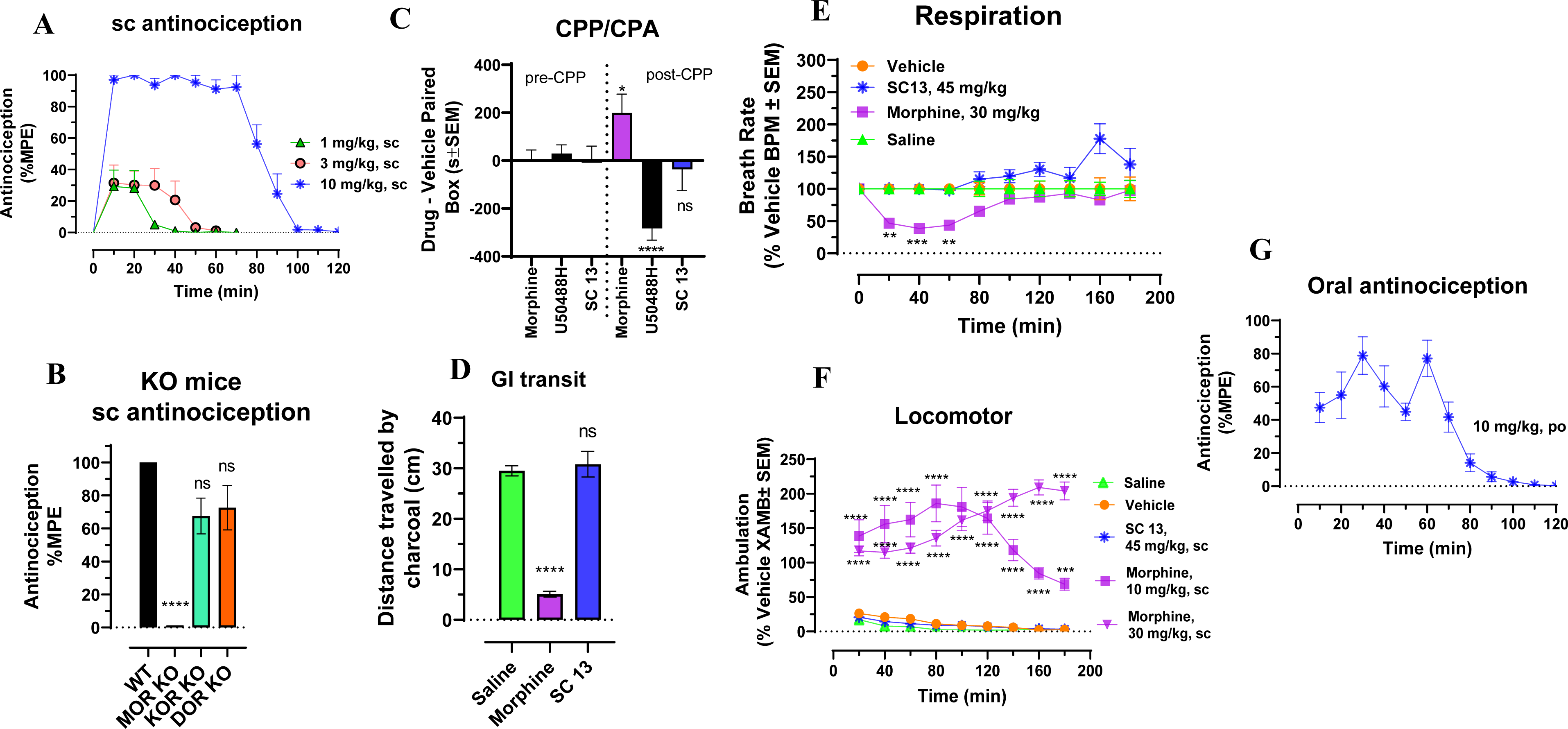
SC13 shows MOR dependent antinociception and lacks abuse potential, constipation, respiratory depression and hyperlocomotion at equianalgesic morphine doses. **A) Antinociception time course**: Groups of C57BL/6J mice were subcutaneously (*sc*) administered **SC13** and antinociception measured using the 55°C tail withdrawal assay. Data are shown as mean % antinociception ± SEM. **(A)** Effect of **SC13** at doses of 1, 3 and 10 mg/kg (*n* = 8 each group) with repeated measures over time. **SC13** showed potent dose dependent antinociception. **B) SC13 antinociception in KO mice**: Antinociception effect of **SC13** (10 mg/kg, *sc*,) was evaluated in group of (n = 8) in WT, MOR KO, KOR KO, and DOR KO mice. Antinociception of **SC13** remained intact in DOR KO (p=0.13) and KOR KO (p=0.058) mice while it was found attenuated in MOR KO. Results for **SC13** were analyzed with one-way ANOVA followed by Tukey’s post-hoc test, F_3,28_=24.07, p<0.0001, ****p<0.0001 relative to WT, ns = p > 0.05 relative to WT. Attenuation of **SC13** antinociception in MOR KO was also significantly greater compared to DOR KO and KOR KO mice (p<0.0001 each; Tukey’s post hoc test). All values are expressed as the mean ± SEM. **C) Conditioned place preference or aversion (CPP/CPA)**: Place conditioning evaluation of **SC13**, morphine, U50,488H, in C57BL/6J mice after *IP* or *sc* administration. Following determination of initial preconditioning preferences, mice were place-conditioned daily for 2 days with **SC13** (15 mg/kg, *sc*; n = 31), U50,488H (30 mg/kg, *IP*; n = 28) or morphine (10 mg/kg, *IP*; n = 18). Mean differences in time spent on the drug-paired side ± SEM are presented. **SC13** does not display significant CPP or CPA compared to the matching preconditioning preference (p<0.05), as determined by unpaired t-test with Welch’s correction. Morphine showed CPP (*p=0.0140) and U50,488H showed CPA (****p<0.0001) and were significantly different from matching preconditioning preference. **D) SC 13 effects on Gastrointestinal Transit**. Mice were administered morphine (10 mg/kg, sc) or **SC13** (15 mg/kg, sc) or saline (0.9%, po), then fed a charcoal meal. After 3h, morphine significantly reduced the distance traveled by the charcoal through the intestines, consistent with the action of a MOR agonist (5.07 ± 0.57 cm, compared to 29.5 ± 1 cm for saline-treated mice; F(2,21) = 81.88, P < 0.0001; one- way ANOVA with Dunnett’s multiple-comparison test. In contrast, compound **SC13** was without significant effect (30.8 ± 2.52 cm). **E) Respiratory rate**: Mice were administered either vehicle (n=12), morphine (30 mg/kg, *sc;* n=12), **SC13** (45 mg/kg, *sc;* n=12) and the breath rates was measured every 20 min for 180 minutes. Morphine administered *sc* caused reduction in the breath rate with respect to saline at 20 min (**p=0.0021), 40 min (***p=0.0003) and 60 min (**p=0.0010) post drug administration. **SC13** (45 mg/kg, *sc*) was not significantly different from vehicle control except at 180 min (****p<0.0001) and 200 min (*p=0.0410) where it showed an increase in breath rates as determined by 2-way ANOVA followed by Dunnett’s multiple-comparison test. **F) Locomotor effect**: Mice were administered either saline (n = 20), vehicle (n=24), morphine (10 and 30 mg/kg, *sc;* n=12 each), and **SC13** (45 mg/kg, *sc;* n=12) and the distance travelled by each group of mice was measured. No significant locomotory effects were observed with **SC13** as determined by 2-way ANOVA followed by Dunnett’s multiple-comparison test in comparsion to vehicle while morphine showed significant hyperlocomotion at every time point compared to saline (p<0.0001). **G) Oral antinociceptive time course**: Groups of C57BL/6J mice were orally (*po*) administered **SC13** at 10 mg/kg and antinociception measured using the 55°C tail withdrawal assay. **SC13** showed antinociception with 80% MPE at peak time point. Data are shown as mean % antinociception ± SEM. Effect of **SC13** at dose 10 mg/kg (*n* = 8) with repeated measures over time.

Notably, as suggested by the coefficient values reported in **Appendix 1-Table 8**, and illustrated in **Appendix 1-Figure 2** by comparing binding modes (**Appendix 1-Figure 2A**) and average SIFts of **SC11-13** with SIFts calculated for morphine (**Appendix 1-Figure 2B**), the **SC11-13** ligands show higher probability of interacting with Y75(1.39), N127(2.63), I144(3.29), H319(7.36), C217(45.50), and W133(23.50), but much lower probability of interacting with H297(6.52), compared to morphine.

### SC13 shows MOR dependent antinociception with reduced adverse effects

The lead MOR-selective agonist **SC13** was characterized in C57BL/6J mice for antinociception, respiratory depression, locomotor effects, inhibition of gastrointestinal transit, and reward or dysphoria (measured using the conditioned place preference or aversion assay (CPP/CPA)).

When administered subcutaneously (sc), **SC13** showed dose-dependent antinociception in mice in the radiant heat 55 °C tail withdrawal assay, with peak effect at 20 min and an ED_50_ (and 95% CI) value of 3.05 (1.75-5.27) mg/kg, sc (**Figure 3A**). Thus, **SC13** potency was similar to that of morphine (ED_50_ = 2.48 (1.57-3.87) mg/kg, sc)^23^, consistent with its roughly comparable Gi-1 potency (EC_50_ = 145 nM compared to morphine (EC_50_ = 51 nM)) as well as in BRET-Nb33 assays (EC_50_ of morphine=584 nM and 1644 and EC_50_ of **SC13**= 12 nM and 730 nM at mMOR and hMOR respectively) for both drugs. Opioid receptor selectivity of **SC13-**mediated antinociception was assessed in transgenic knock-out (KO) mice lacking MOR, KOR or DOR. **SC13** antinociception was significantly reduced in MOR KO mice (**Figure 3B**). DOR KO did not produce significant differences in effect from WT mice, and while KOR contributions were trending towards significance, they did not reach statistical threshold. Blockage of **SC13** antinociception in MOR KO was significantly greater compared to DOR KO and KOR KO mice supporting the conclusion that **SC13** antinociception was predominantly MOR-mediated. The results were also consistent with **SC13** selectivity seen in Gi-1 BRET assays.

At doses 5-fold higher than their ED_50_ antinociceptive values, **SC13** (15 mg/kg, sc) showed no signs of CPP or CPA whereas morphine (10 mg/kg, *IP*) showed CPP and U50,488H showed CPA, as expected (**Figure 3C**). In GI transit assays tested at ED_80_ antinociceptive doses, morphine inhibited gastrointestinal passage, while the effects of **SC13** and saline were indistinguishable from each other (**Figure 3D**).

Compounds were next evaluated for respiratory depression and hyperlocomotion in mice using the computer-controlled Comprehensive Lab Animal Monitoring System (CLAMS) assay^26^. At a dose 15-fold higher than the antinociceptive ED_50_ value, **SC13** showed no statistically significant respiratory effects, whereas morphine at an equivalent dosage (30 mg/kg, sc) showed significant respiratory depression for 60 min after administration (**Figure 3E**). Similarly, **SC13** showed no hyperlocomotion at a dose 15-fold higher than the antinociceptive ED_50_ value, in contrast to the prototypic MOR agonist morphine, which showed hyperlocomotion effects at doses 5-fold and 15-fold higher than its antinociceptive ED_50_ value (**Figure 3F**).

Oral administration of **SC13** (10 mg/kg, po) also showed antinociceptive efficacy (E_max_ = 80% MPE at 30 min) nearly equivalent to the efficacy of subcutaneous **SC13** at the same dose (E_max_=100% at 20 min), suggesting possibly good plasma exposure through the oral route. (**Figure 3G**). The antinociceptive time courses observed following administration by each route were also similar. The results are consistent with the good oral activity usually seen with the mitragynine template^19, 42^ and reported metabolic stability of this template.^19^ Overall, the MOR partial agonist, **SC13** with an efficacy of ∼10% (BRET-Nb33 assays, **Appendix 1-Figure 1J**) in murine MOR showed equi-efficacious antinociception compared to morphine with 70% efficacy ((BRET-Nb33 assays, **Appendix 1-Figure 1J**) while showing greatly attenuated opioid-induced adverse effects in mice.

## DISCUSSION

Opioids and their activation of opioid receptors continue to be investigated as treatments of acute to moderate pain despite their numerous and often serious adverse effects. In recent years, biased agonism has been proposed as an avenue to dissociate respiratory depression from analgesia.^43–45^ However, recent studies have raised concerns about this hypothesis.^44, 46, 47^ Mice lacking β-arrestin2 were reported to retain respiratory depression mediated by morphine,^48^ and mice with MOR C-tail mutations that inhibit arrestin recruitment still show respiratory depression as well as tolerance^49^ in contrast to previous reports^50^.

Extending these concerns, we had previously reported the kratom alkaloids mitragynine, 7OH and mitragynine pseudoindoxyl to be G-protein biased agonists.^17, 18, 22^ However, recent reports with other putative G-protein biased agonists such as SR17018, PZM21 and TRV130 have suggested that these ligands are in fact MOR partial agonists with low intrinsic efficacy compared to DAMGO when assessed in a less amplified G-protein signaling system.^5^

Here we used the mitragynine template to test this low efficacy partial agonism hypothesis, and whether such an approach can lead to MOR agonists with reduced side effect liability but maintained analgesia. We developed a SAR based on the aromatic ring of mitragynine, 7OH template and identified three C9-diversified analogs **SC11, SC12** and **SC13.**

In amplified cAMP assays, our analogs showed full agonism at MOR compared to DAMGO. Similar observations of cAMP measurements greatly overestimating efficacy in the presence of receptor reserve have been reported previously ^51^ and are consistent with receptor theory.^52^ Using a less amplified TRUPATH assay, we find that the three lead analogs have less efficacy relative to DAMGO, fentanyl and morphine but higher efficacy than buprenorphine, a well characterized partial agonist at MOR.^53^

The lead analogs, **SC11-13** also showed arrestin recruitment with poor potency when assessed using TANGO (an assay with amplified signaling), but this β-arrestin2 recruitment activity was significantly reduced or altogether absent when quantified in the less amplified DiscoverX Pathhunter or BRET-based assays.

Typical opioid receptor functional assays that utilize cAMP and [^35^S]GTP*γ*S often fail to account for simultaneous signaling through various G*α* subunits.^54^ It is difficult to recapitulate the complexity of *in vivo* signaling due to cell line limitations, namely, the differential expression of specific G*α* subunits in various cell types. For example, CHO and HEK cell lines show differential expression of Gz and Gαo subtypes.^33^ The BRET based TRUPATH assay enabled us to study activity of each of the Gα-subtypes in isolation.^32^ We determined that **SC11-13** show lower efficacy than DAMGO, morphine, or fentanyl at each Gi/o subtype. At GoA and GoB, buprenorphine (E_max_=65-66%) and **SC11-13** (E_max_=63-75%) had comparable intrinsic efficacy, which is of interest since the most abundant G*α* subunit in the brain is GoA._55_ Similarly at Gz too, buprenorphine (E_max_=81%) and our synthetic analogs **SC11-13** (E_max_=79-86%) had similar efficacies.

In mice, **SC13** was equipotent to morphine in antinociception assays. The role of Gz in opioid induced antinociception is poorly understood, although Gz knock-out (KO) mice have reduced opioid antinociception in a tail withdrawal test similar to the one used here,^56^ and DAMGO preferentially signals through Gz over Gi-2 in periaqueductal grey membranes.^57^ The role these Gα subtypes play in *in vivo* responses to **SC11-13** is uncertain at this point; however the overall Gα-subtype efficacy profile does appear similar to the well characterized MOR partial agonist buprenorphine.

The partial agonism of **SC11-13** was confirmed in a BRET-based Nb33 recruitment, an assay which has been shown to accurately reflect efficacy without signal amplification^5^. In these assays, using either hMOR or mMOR, **SC11-13** was found to have similar efficacy to buprenorphine. While the putatively biased MOR ligands SR17018, TRV130 and PZM21 were not evaluated in our study, we infer that **SC11-13** may have efficacy similar to SR17018 (20%) but lower than either TRV130 (42%) or PZM21 (38%).^5^ Similarly, **SC11-13** were found to have lower intrinsic efficacy than DAMGO and morphine in VTA synaptic effects, corroborating our cell line-based findings in an endogenous system with physiologically relevant levels of receptor reserve. Behaviorally, **SC13** showed MOR-dependent antinociception and potency similar to morphine while showing none of the adverse effects associated with morphine at equianalgesic equivalent doses. This pattern is reminiscent of that of buprenorphine, which is known to show a ceiling effect in respiratory depression^58^ and is generally considered a safer analgesic^59^ although it still shows hyperlocomotion^60^, constipation^60^ and reward-like behavior in rodents.^61^ Of note, it is not yet clear if the preferable properties of buprenorphine result solely from its partial agonism at MOR^5, 62^ or because of its additional actions such as DOR^60^ and KOR^60^ antagonism or weak NOP agonism^63^. Buprenorphine’s pharmacology is further complicated by its metabolism to norbuprenorphine^64, 65^ (a lower potency but much higher efficacy metabolite) as well as other active metabolites such as buprenorphine 3-glucuronide^66^. Furthermore, the oral activity of buprenorphine is limited due to its metabolism to norbuprenorphine^67^, unlike **SC13** which under tested conditions in mice is orally as active as when given subcutaneously. Together, the present results suggest additional benefits of **SC13** over buprenorphine, while also validating the further investigation of MOR-selective partial agonists as analgesics with fewer liabilities.

In summary, using the mitragynine template and unamplified signaling assays we identified partial MOR agonists that appear to functionally dissociate MOR-dependent analgesia from locomotor and respiratory depression. While additional mechanisms extending beyond MOR and Gα signaling cannot be ruled out, our studies corroborate findings by Gillis et al^5^ suggesting that low G protein efficacy at MOR may lead to a favorable therapeutic window of new opioids.

## ACKNOWLEDGEMENTS

SM and DS are supported by funds from NIH grants DA045884, DA046487. SM is supported by DA048379. JAJ is supported by a grant from the Hope for Depression Research Foundation. MF is supported by NIH grants DA034049 and DA045884. Computations were run on resources available through the Office of Research Infrastructure of the National Institutes of Health under award numbers S10OD018522 and S10OD026880 (to the Icahn School of Medicine at Mount Sinai), as well as the Extreme Science and Engineering Discovery Environment under MCB080077 (to MF), which is supported by National Science Foundation grant number ACI- 1548562. RVR is supported by NIH grants AA026949, DA045897, AA025368. EBM is supported by funds from the State of California for medical research on alcohol and substance abuse through the University of California, San Francisco.

## METHODS

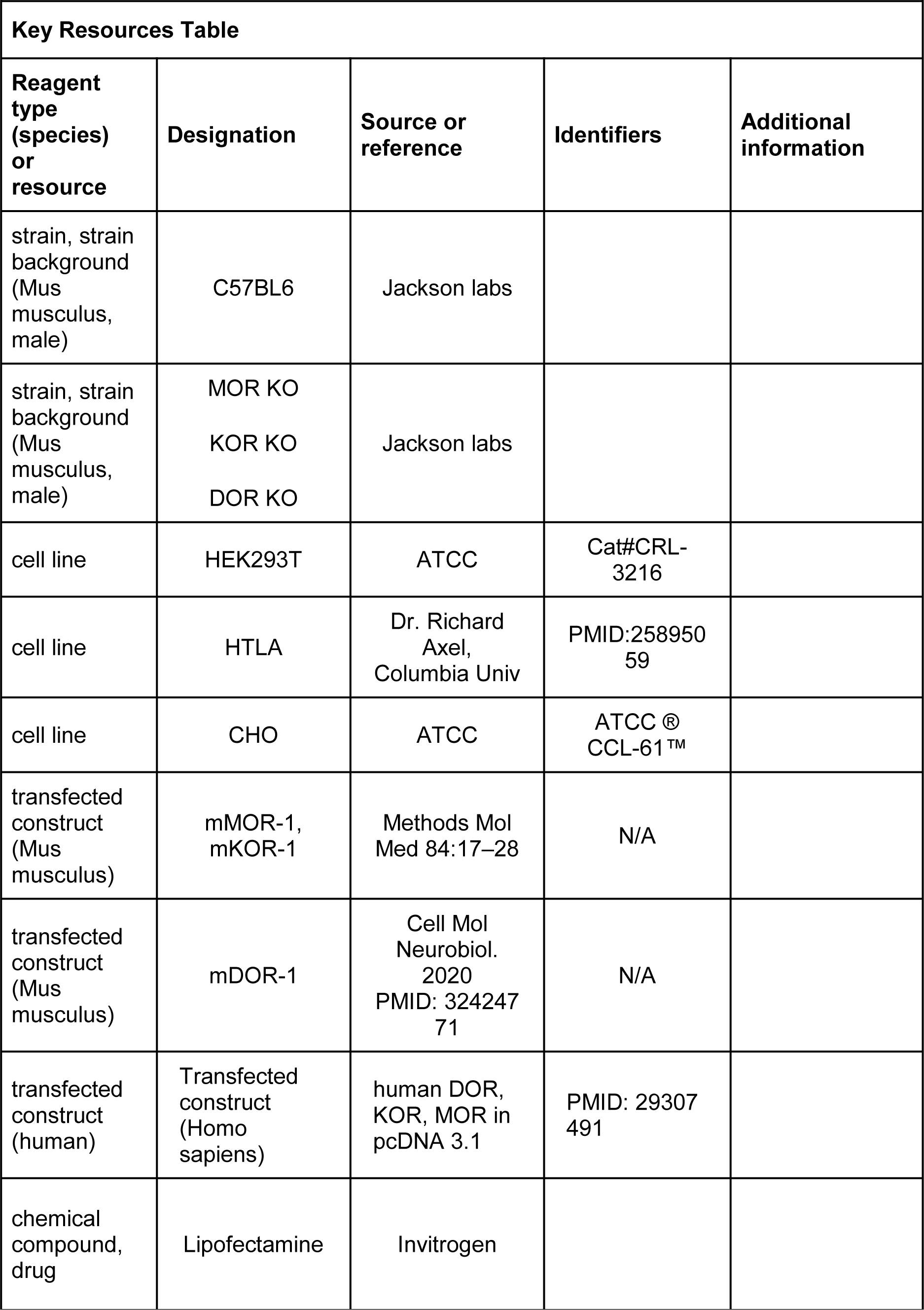
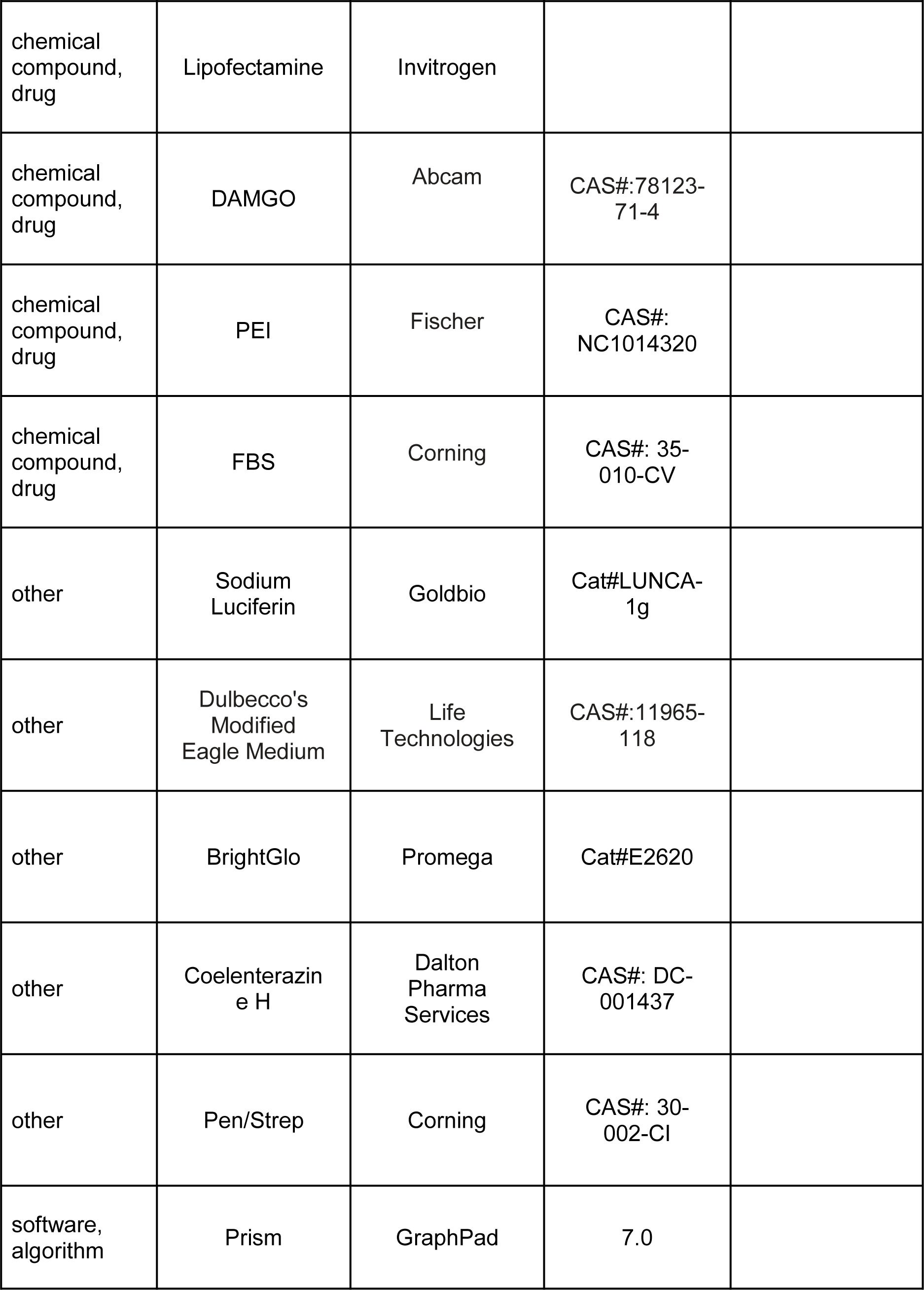
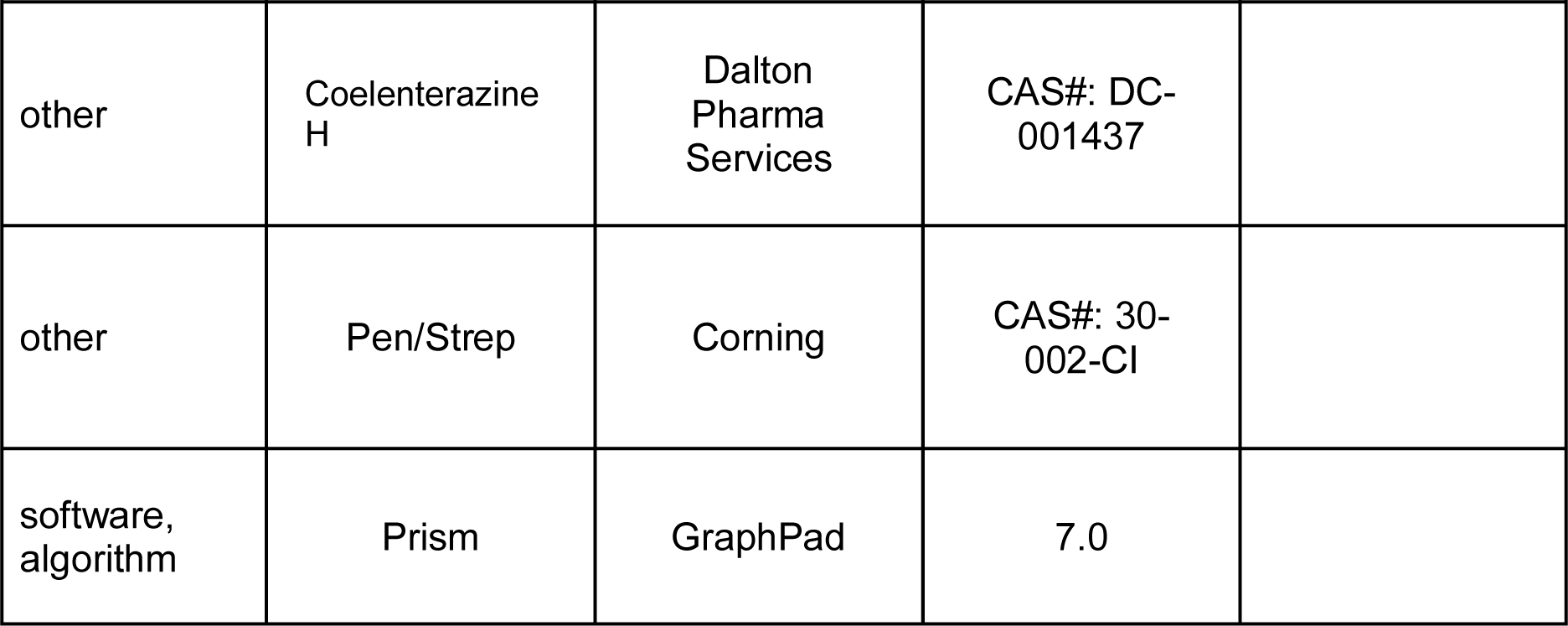

### Drugs and Chemicals

Opiates were provided by the Research Technology Branch of the National Institute on Drug Abuse (Rockville, MD). Selective opioid antagonists were purchased from Tocris Bioscience. Miscellaneous chemicals and buffers were purchased from Sigma-Aldrich. Kratom “Red Indonesian Micro Powder” was purchased from Moon Kratom (Austin, TX).

### Chemistry

All chemicals were purchased from Sigma-Aldrich Chemicals and used without further purification. Reactions were carried out in flame-dried reaction flasks under Ar. Reaction mixtures were purified by silica flash chromatography on E. Merck 230−400 mesh silica gel 60 using a Teledyne ISCO CombiFlash Rf instrument with UV detection at 280 and 254 nm. RediSep R_f_ silica gel normal phase columns were used. The yields reported are isolated yields. NMR spectra were recorded on a Varian 400/500 MHz NMR spectrometer. NMR spectra were processed with MestReNova software. The chemical shifts were reported as δ ppm relative to TMS using residual solvent peak as the reference unless otherwise noted (CDCl3 ^1^H: 7.26, ^13^C: 77.3). Peak multiplicity is reported as follows: s, singlet; d, doublet; t, triplet; q, quartet; m, multiplet. Coupling constants (*J*) are expressed in Hz. High resolution mass spectra were obtained on a Bruker Daltonics 10 Tesla Apex Qe Fourier Transform Ion Cyclotron Resonance-Mass Spectrometer by electrospray ionization (ESI). Accurate masses are reported for the molecular ion [M + H]^+^.

### In vitro pharmacology assays

#### cAMP and TANGO.^34^

To measure Glo-sensor G_αi_-mediated cAMP inhibition, HEK 293T (ATCC CRL-11268) cells were co-transfected with human opioid receptor (hMOR, hKOR and hDOR) along with a luciferase-based cAMP biosensor and the assay was performed as reported previously.^34^ Next, arrestin recruitment Tango assay was carried out using HTLA cells expressing TEV fused-β- Arrestin2 were transfected with human opioid receptors (hMOR, hKOR or hDOR) as the Tango construct by following previously reported protocols.^34^

#### BRET2 assays^32^

Cells were plated either in 6-well dishes at a density of 700,000–800,000 cells per well, or 10 cm dishes at 7–8 million cells per dish. Cells were transfected 2–4 h later, using a 1:1:1:1 DNA ratio of receptor:Gα-RLuc8:Gβ:Gγ-GFP2 (100 ng per construct for 6-well dishes, 750 ng per construct for 10 cm dishes), except for the Gγ-GFP2 screen, where an ethanol co-precipitated mixture of Gβ1–4 was used at twice its normal ratio (1:1:2:1). Transit 2020 (Mirus Biosciences) was used to complex the DNA at a ratio of 3 µl Transit per µg DNA, in OptiMEM (Gibco-ThermoFisher) at a concentration of 10 ng DNA per µl OptiMEM. The next day, cells were harvested from the plate using Versene (0.1 M PBS + 0.5 mM EDTA, pH 7.4) and plated in poly-D-lysine-coated white, clear-bottom 96-well assay plates (Greiner Bio-One) at a density of 30,000–50,000 cells per well. One day after plating in 96-well assay plates, white backings (PerkinElmer) were applied to the plate bottoms, and growth medium was carefully aspirated and replaced immediately with 60 µl of assay buffer (1× Hank’s balanced salt solution (HBSS) + 20 mM HEPES, pH 7.4), followed by a 10 µl addition of freshly prepared 50 µM coelenterazine 400a (Nanolight Technologies). After a 5 min equilibration period, cells were treated with 30 µl of drug for an additional 5 min. Plates were then read in an LB940 Mithras plate reader (Berthold Technologies) with 395 nm (RLuc8-coelenterazine 400a) and 510 nm (GFP2) emission filters, at integration times of 1 s per well. Plates were read serially six times, and measurements from the sixth read were used in all analyses. BRET2 ratios were computed as the ratio of the GFP2 emission to RLuc8 emission.

### BRET-Based Nb33 Recruitment Assays

Experiments were performed as described previously.^68^ Briefly, transfected cells were dissociated and resuspended in phosphate-buffered saline. Cells were added to a black-framed, white well 96-well plate (no. 60050; Perkin Elmer; Waltham, MA, USA). At time zero, the luciferase substrate coelenterazine H (5 μM) was added to each well. Ligands were added after 5 min, then BRET signal was measured 10 min later. BRET measurements were performed using a PHERAstar FS plate reader (BMG Labtech, Cary, NC, USA). The BRET signal was calculated as the ratio of the light emitted by the mVenus acceptor (510–540 nm) over the light emitted by the NanoLuc donor (475 nm). Dose–response curves were fit using a three-parameter logistic equation in GraphPad Prism 8 (Graphpad Software, La Jolla, CA, USA). All experiments were repeated in at least three independent trials each with triplicate determinations.

#### Materials

HEK-293T cells were obtained from the American Type Culture Collection (Rockville, MD, USA) and were cultured in a 5% CO_2_ atmosphere at 37 °C in Dulbecco’s Modified Eagle Medium (DMEM, high glucose, #11965; Life Technologies; Grand Island, NY, USA) supplemented with 10% Fetal Bovine Serum (#35-010-CV, Corning, Corning, NY, USA) and 100 IU ml^−1^ penicillin and 100 μg ml^−1^ streptomycin (#30-002-CI; Corning, Corning, NY, USA).

The following chemicals were used without further modification: [D-Ala^2^, N-Me-Phe^4^, Gly^5^-ol]- Enkephalin (DAMGO; #78123-71-4,Abcam,Cambridge, United Kingdom), Buprenorphine hydrochloride (#B9275, Sigma-Aldrich,St. Louis, MO, USA), Morphine sulfate (#M1167, Spectrum Chemicals,New Brunswick, NJ, USA), Coelenterazine H (#DC-001437, Dalton Pharma Services, Toronto, ON, Canada), Polyethylenimine (PEI; #NC1014320, Polysciences, Warrington, PA, USA).

#### DNA Constructs

The expression vector coding for mouse MOR tagged at the C-terminus with Nanoluc (mMOR-nluc) by a Gly-Ser linker was constructed using standard techniques in molecular biology and confirmed by DNA sequencing (Psomagen, Brooklyn, NY, USA). Briefly, two DNA inserts were PCR amplified, one coding for mMOR with an N terminal signal peptide followed by a FLAG tag, and the other coding for NanoLuc. The two inserts were joined by PCR amplification and the resulting insert coding mMOR-nluc was cloned into the *Hind III* and *Xho I* sites of pcDNA3.1 (+) (#V79020, ThermoFisher Scientific, Waltham, MA, USA). The plasmid coding for human MOR-nanoluc (hMOR-nluc) was a gift from Dr. Nevin Lambert at the Medical College of Georgia. The plasmid coding for the nanobody-33-Venus (Nb-33) construct^5^ was a gift from Dr. Meritxell Canals at the University of Nottingham.

#### Transfection

A total of 5 μg of cDNA was transiently transfected into HEK-293T cells (2 × 10^6^ cells per plate) in 10 cm dishes (1 μg receptor-nluc, and 4 μg Nb-33-Venus), using PEI in a 6:1 ratio (diluted in DMEM). Cells were maintained in the HEK-293T media described above. Experiments were performed 48 hours after transfection.

### Pathhunter assays

β-arrestin recruitment assays were performed as previously described.^69^ In brief, CHO-K1-human µOR cells (DiscoverX) were grown to confluency and seeded at a density of 2500 cells in a low-volume 384 well plate (10 µL per well). After incubating overnight at 37°C with 5% CO2, a 5X dilution series of compounds prepared in opti-MEM was added (2.5 µL per well) and incubated at 37°C for an additional 90 minutes. PathHunter detection reagent (DiscoverX) was prepared according to the manufacturer’s protocol and added (6 µL per well). Following a 60-minute, room temperature incubation in the dark, the chemiluminescence signal was measured using a FlexStation3 plate reader.

### Competitive radioligand binding assay

Membrane isolation and binding assays were performed as previously described.^70^ In brief, membranes were isolated from CHO cells stably expressing the µOR (DiscoverX). To harvest membranes, cells were dislodged from a T75 flask and pelleted via centrifugation at 1300 rpm for 5 minutes at 20 °C (Eppendorf 5804R). The supernatant was aspirated, and the cell pellet was resuspended in assay buffer (50 mM Tris HCl, 10 mM MgCl_2_, 1 mM EDTA, pH 7.4) and thoroughly sonicated (Qsonica XL-2000, level 3.) Membranes were isolated from the resulting suspension via ultracentrifugation at 20,000 rpm for 30 minutes at 4 °C (Optima L-100 XP Ultracentrifuge, SW 41 Ti, 41,000 rpm rotor). The supernatant was aspirated, and the resulting membrane pellets were resuspended in assay buffer on ice by thorough sonication, pushed through a 28-gauge needle, and stored in 1 mL aliquots at -80 °C until day of binding assay. Each T75 flask yielded approximately 1, 1 mL aliquot. On assay days, a 4X dilution series of compounds made in assay buffer was added to a 96 well plate (50 µL per well, added in duplicate.) Tritiated radioligand ([3H]DAMGO for MOR) diluted in assay buffer was added to the 96 well plate (50 µL per well) at a concentration near the EC80 value for the receptor: 2.325 nM [3H]DAMGO. Next, a membrane aliquot was thawed on ice, diluted in assay buffer (1:10, approximate protein concentration of 70 µg/mL) followed by thorough sonication, and the resulting membrane suspension was added to the plate (100 µL per well, approximately 7 µg protein). After adding the membrane suspension, the plate contents were incubated at room temperature for 90 minutes. The membrane mixture was then filtered over a 0.3% PEI pre-treated GF-B/C plate (#6005174, Perkin Elmer, Waltham, MA, USA) with a cell harvester system. After the GF-B/C plate was dried overnight, scintillation fluid was added (50 µL per well, Ultimagold uLLT) and radioactivity was measured using a scintillation counter (Hewlett Packard TopCount NXT). For the competitive binding assays, all data were analyzed with GraphPad 8 (GraphPad Prism software, La Jolla, CA). Both assays were run in duplicate in a minimum of three, independent assays. Data from each independent assay were normalized to a positive control and then all independent assays were averaged and compiled into a composite Figure. Data is presented as means ± SEM.

### EPhys assays

#### Electrophysiology Animals

Eight male Sprague-Dawley rats were used for whole cell electrophysiology recordings, procedures conducted in strict accordance with the recommendations of the National Institutes Health (NIH) in the Guide for the Care and Use of Laboratory Animals. Research protocols were approved by the Institutional Animal Care and Use Committee (University of California at San Francisco, CA), approval ID AN183735-01B.

#### Slice preparation and *ex vivo* whole cell electrophysiology^71^

Rats were anesthetized with isoflurane and their brains were removed. Horizontal brain slices (200 mm thick) containing the VTA were prepared using a vibratome (Leica Microsystems). Slices were submerged in artificial CSF solution containing (in mM): 126 NaCl, 2.5 KCl, 1.2 MgCl, 1.4 NaH_2_PO_4_, 2.5 CaCl_2_, 25 NaHCO_3_, and 11 glucose saturated with 95% O_2_ – 5% CO_2_ and allowed to recover at 33°C for at least 1 h. Individual slices were visualized under a Zeiss AxioExaminer.D1 with differential interference contrast, Dodt, and near infrared optics using a monochrome Axiocam 506 or under a Zeiss Axioskop FS 2 plus with differential interference contrast optics and infrared illumination equipped with a Zeiss Axiocam MRm (Zeiss International). Whole-cell patch-clamp recordings were made at 33°C using 2.5–5 MW pipettes containing (in mM) 128 KCl, 20 NaCl, 1 MgCl_2_, 1 EGTA, 0.3 CaCl_2_, 10 HEPES, 2 MgATP, and 0.3 Na_3_GTP (pH 7.2, osmolarity adjusted to 275). Signals were amplified using an IPA amplifier with SutterPatch software (Sutter Instrument) filtered at 1 kHz and collected at 10 kHz or using an Axopatch 1-D (Molecular Devices), filtered at 2 kHz, and collected at 20 kHz using custom written procedures for IGOR Pro (Wavemetrics). Cells were recorded in voltage-clamp mode (V -70 mV). Series resistance and input resistance were sampled throughout the experiment with 4 mV, 200 ms hyperpolarizing steps. GABA_A_ receptor mediated inhibitory postsynaptic potentials (IPSCs) were pharmacologically isolated with 6,7-dinitroquinoxaline-2,3(1H,4H)-dione (DNQX: 10 mM). Stimulating electrodes were placed 80–250 mm anterior or posterior to the soma of the recorded neuron. To measure drug effects on evoked IPSCs, paired pulses (50-ms interval) were delivered once every 10 s. The IPSC amplitude was calculated by comparing the peak PSC voltage to a 2 ms interval just before stimulation. All drugs were bath applied. Drug effects were quantified by comparing the mean evoked IPSC amplitude during the 4 min of baseline just preceding drug application and the mean response amplitudes during minutes 4–7 of drug application.

#### Mice

C57BL/6J mice (20–32 g each) were obtained from Jackson Laboratories (Bar Harbor, ME). MOR KO, KOR KO and DOR KO were bred in the laboratory of Dr. McLaughlin at University of Florida. All mice used throughout the manuscript were opioid naïve. All mice were maintained on a 12-hour light/dark cycle with Purina rodent chow and water available ad libitum, and housed in groups of five until testing. All animal studies were preapproved by the Institutional Animal Care and Use Committees of the University of Florida, in accordance with the 2002 National Institutes of Health Guide for the Care and Use of Laboratory Animals

#### Antinociception

The 55 °C warm-water tail-withdrawal assay was conducted in C57BL/6J mice as a measure of acute thermal antinociception as described previously.^18^ Briefly, each mouse was tested for baseline tail-withdrawal latency prior to drug administration. Following drug administration, the latency for each mouse to withdraw the tail was measured every 10 minutes until latency returned to the baseline value. A maximum response time of 15 seconds was utilized to prevent tissue damage. If the mouse failed to display a tail-withdrawal response within 15 seconds, the tail was removed from the water and the animal was assigned a maximal antinociceptive score of 100%. Data are reported as percent antinociception, calculated by the equation: % antinociception = 100 x [(test latency - baseline latency)/ (15 - baseline latency)]. This was utilized to account for innate variability between mice. Compounds were administered subcutaneously (sc) or orally (po) and the analgesic action of compounds was assessed at the peak effect.

#### Respiratory Depression and Locomotor Effects Assessment

Respiration rates and spontaneous ambulation rates were monitored using the automated, computer-controlled Comprehensive Lab Animal Monitoring System (CLAMS) (Columbus Instruments, Columbus, OH) as described previously.^26^ Freely moving mice were habituated in closed, sealed individual apparatus cages (23.5 cm × 11/5 cm × 13 cm) for 60 min before testing. To start testing, mice were administered (sc) drug or vehicle and 5 min later confined to the CLAMS testing cages for 120 min. Using a pressure transducer built into the sealed CLAMS cage, the respiration rate (breaths/min) of each occupant mouse was measured. Infrared beams located in the floor measured locomotion as ambulations, from the number of sequential breaks of adjacent beams. Data are expressed as percent of vehicle control response.

#### Conditioned Place Preference and Aversion

Mice were conditioned with a counterbalanced place conditioning paradigm using similar timing as detailed previously. A group of mice (n = 18-24) were habituated to freely explore both sides of a two-compartment apparatus for 3 hours each for 2 days prior testing. The amount of time subjects spent in each of three compartments was measured over a 20 min testing period. Prior to place conditioning, the animals did not demonstrate significant differences in their time spent exploring the left vs right compartments. During each of the next 2 days, mice were administered vehicle (0.9% saline) and consistently confined in a randomly assigned outer compartment for 20 min, half of each group in the right chamber, half in the left chamber. Four hours later, mice were administered drugs morphine (10 mg/kg/d, *IP*), U50,488h (30 mg/kg/d, *IP*), **SC 13** (15 mg/kg/d, sc) and were confined to the opposite compartment for 20 min. Conditioned place preference data are presented as the difference in time spent in drug- and vehicle-associated chambers and were analyzed via repeated measures two-way ANOVA with the difference in time spent on the treatment- vs vehicle-associated side as the dependent measure and conditioning status as the between groups factor. Where appropriate, Tukey’s HSD or Sidak’s multiple comparison post hoc tests were used to assess group differences. Effects were considered significant when p < 0.05. All effects are expressed as mean ± SEM.

#### Assessment of Gastrointestinal Transit

C57BL/6J mice (8 per drug treatment) were administered morphine (10 mg/kg, sc), or saline (0.9%, sc) or **SC13** (15 mg/kg, sc) 20 min prior to oral gavage with 0.3 mL of a 5% aqueous solution of charcoal meal. After 3 h, mice were euthanized and the intestines removed. The progression of charcoal through the intestines was measured as distance traveled from the jejunum to the cecum as utilized elsewhere.^72^

### Computational studies

#### Molecular Docking

The crystal structure of active murine MOR bound to BU72 (PDB id: 5C1MF) was prepared for molecular docking of **SC1, SC3, SC4, SC11, SC12, SC13, 11-F**, morphine, and buprenorphine, using the protocol we recently reported in the literature for docking and simulations of kratom alkaloids, including mitragynine and 7OH mitragynine.^73^ Molecular docking of morphine and buprenorphine was achieved by overlapping core heavy atoms onto the co-crystal compound BU72. In contrast, **SC1, SC3, SC4, SC11, SC12, SC13**, and **11-F** were aligned onto mitragynine and 7OH mitragynine binding poses that had been previously obtained^73^ using the *Binding Pose Metadynamics* module in Schrödinger suite 2019-2^74^ for metadynamics rescoring of initial docking poses obtained with DOCK6.9.

#### Molecular Dynamics Simulations

Unbiased MD simulations of ligand-MOR complexes embedded in a POPC bilayer and solvated in a 10×10×10 Å^3^ orthorhombic box of simple point charge (SPC) water molecules and 0.15 M NaCl buffer in each dimension, were carried out using the OPLS3e force-field and the Desmond software within the Schrödinger suite 2019-2.^39^ Systems were neutralized with chloride ions using the *System Builder* function and missing dihedral parameters of the ligands were generated using the *Force Field Builder* in the Schrödinger suite. The same MD simulation parameters and protocol used in our previous work on 7OH mitragynine and mitragynine^39^ were used here. MD production runs consisted of four independent simulations of 250 ns each for each ligand-MOR complex, for a total of 9 μs new simulation data added to the previously published 2 μs simulation data collected for 7OH-mitragynine-MOR and mitragynine-MOR complexes.^39^ Highly populated conformations of each ligand at MOR were obtained using the affinity propagation clustering algorithm described by Fray and Dueck^75^ and implemented in the Schrödinger’s *trj_cluster.py* script. Specifically, 500 snapshots of each ligand-MOR MD simulation trajectory with a stride of 2 ns were superimposed to a reference frame using the protein heavy atoms within 8 Å of the ligand prior to clustering. Pairwise root mean square deviation (RMSD) values of the same selected group of atoms were used as input for *trj_cluster.py*, which yielded 39, 39, 36, 39, 41, 28, 41, 46, 36, 41, and 43 clusters for mitragynine, 7OH- mitragynine, morphine, buprenorphine, **SC1, SC3, SC4, SC11, SC12, SC13**, and **11-F** respectively. The top populated cluster in each case accounted for 4.58%, 5.38%, 8.96%, 6.57%, 6.57%, 8.37%, 6.97%, 4.78%, 10.36%, 5.58% and 6.17% of the assessed simulation frames.

#### Structural Interaction Fingerprint (SIFt) Analysis

An in-house python script was used to generate 9-bit representations of ligand-receptor interactions formed by both backbone and sidechain atoms, and including hydrogen-bond interactions with the protein as hydrogen-bond donor (Hbond_proD) or hydrogen-bond acceptor (Hbond_proA), electrostatic interactions with positively (Elec_ProP) or negatively charged (Elec_ProN) residues, apolar interactions (carbon-carbon atoms in contact), face-to-face (Aro_F2F) and edge-to-face (Aro_E2F) aromatic interactions, as well as 1-water mediated H-bond (Hbond_1wat), and 2-water mediated H-bond (Hbond_2wat). Apolar interactions were cut at 4.5 Å whereas a cutoff of 4 Å was considered to define aromatic and electrostatic interactions. A two-state Markov model that samples the transition matrix posterior distribution using standard Dirichlet priors for the transition probabilities as described by Noé et al was used to calculate the probability of each ligand-MOR interaction formed during MD simulations.^76^ Calculated average SIFt probabilities for each ligand are listed in **Appendix 1-Table 6.**

#### Logistic Regression Models Based on SIFTs

We modeled the negative logarithm of the G protein efficacy *E*_*max*_ for each ligand *k* as a function of the probability of ligands establishing up to three interactions *p*_*i*_(*k*) in the binding pocket according to the equation:

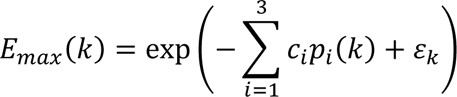

where *ε*_*k*_ is a normally distributed error term and *c*_*i*_ are scalar coefficients. According to this equation, high probability of establishing an interaction whose coefficient *c*_*i*_ is negative results in enhancing the efficacy of the ligand, while the formation of an interaction whose coefficient *c*_*i*_ is positive reduces the ligand’s efficacy. The models were estimated in a Bayesian framework using the STAN engine^77^ for all possible combinations of three interactions in the binding pocket. The accuracy and robustness of the model was assessed by calculating the R^2^ on the full training dataset (11 ligands), as well as the RMSE in a LOO validation. The best 8 performing models on the experimental data were those in the top quartile of R^2^ validation on the full training set and the lowest LOO-RMSE validation (red dots in **Appendix 1-Figure 3**). To summarize the effect of each of the interactions identified by these 8 models, we report the average coefficients as well as the number of times the interactions appear in the top 8 models in **Appendix 1-Table 8**.

### Chemistry

#### Synthesis of C9 analogs

Kratom “Red Indonesian Micro Powder” was purchased from Moon Kratom (Austin, TX). Mitragynine (**1)** was extracted from dry kratom powder using a modified protocol reported by Varadi et al.^18^ 500 g of kratom powder was used to isolate 4.5 g of mitragynine along with other alkaloids. **1** was converted to 9-hydroxymitragynine using AlCl_3_ and ethanethiol in DCM using literature repoeted procedure.^16^ This hydroxy compound was converted to its triflate (**4**) using N-Phenyl-*bis*(trifluoromethanesulfonimide) and Et_3_N in DCM, which was subsequently used as the precursor for further reactions.

As shown in Figure 1B, 9-phenyl mitragynine (**SC3**) was synthesized in 65% yield using palladium-catalyzed Suzuki coupling reaction of triflate **4** with phenylboronic acid. **SC3** was then converted to the corresponding 7OH derivative **SC11** in 33% yield using oxone® and aqueous NaHCO3. The synthesis of 9-3’-furanyl mitragynine (**SC1)** was accomplished by a similar palladium-catalyzed reaction of triflate **4** with 3-furanylboronic acid. Alcohol **SC13** was obtained via oxidation of **SC1** using oxone® and aqueous NaHCO_3_. To install the methyl group at C9 we used DABAL-Me_3_ as methyl donor. Palladium catalyzed coupling reaction of triflate **4** with DABAL-Me_3_ in presence of XPhos afforded 9-methyl mitragynine (**SC4**) in 68% isolated yield. Oxidation of compound **SC4** using oxone® and aqueous NaHCO_3_ resulted in hydroxide **SC12**.

#### Synthesis of C10 analogs

To have access to the C10 position of the mitragynine scaffold, we incorporated bromide selectively at C10 position using Takayama’s protocol.^31^ Mitragynine (**1**) was converted to mitragynine-ethylene glycol adduct using PIFA and ethylene glycol (Figure 1C) followed by bromination with NBS in DMF gave 10 bromo derivative **5**^31^ in 74% yield along with 24% of 12 bromo derivative. In this adduct the indole’s double bond is temporarily masked by an ethylene glycol group. The deprotection of **5** to 10 bromo mitragynine (**6**)^31^ was carried out by a mild reductive condition using NaBH3CN. ^1^H NMR of **6** was in good agreement with the literature reported value.^31^ ^1^H NMR (500 MHz, Chloroform-*d*) δ 7.86 (s, 1H), 7.43 (s, 1H), 7.18 (d, *J* = 8.5 Hz, 1H), 6.94 (d, *J* = 8.5 Hz, 1H), 3.92 (s, 3H), 3.74 (s, 3H), 3.71 (s, 3H), 3.19 – 3.08 (m, 2H), 3.07 – 3.01 (m, 2H), 3.00 – 2.90 (m, 2H), 2.59 – 2.43 (m, 3H), 1.84 – 1.74 (m, 2H), 1.66 – 1.62 (m, 1H), 1.24 – 1.18 (m, 1H), 0.87 (t, *J* = 7.4 Hz, 3H).

10-bromo mitragynine **6**, was submitted to different coupling reactions to furnish analogs of C10 mitragynine. 10-phenyl mitragynine (**SC21**) was synthesized in 71% yield using palladium-catalyzed coupling reaction of bromide **6** with phenylboronic acid. **SC21** was then treated with oxone® and aqueous NaHCO_3_ to furnish the corresponding 7OH derivative **SC31** in 31% yield. The synthesis of 10-3’-furyl mitragynine (**SC22)** was accomplished by a similar palladium-catalyzed reaction of bromide **6** with 3-furanylboronic acid. Treatment of **SC22** with oxone® and aqueous NaHCO_3_ produced alcohol **SC32**. The methyl group at C10 position was introduced by DABAL-Me_3_. Palladium catalyzed coupling reaction of bromide **6** with DABAL-Me_3_ in presence of XPhos afforded 10-methyl mitragynine (**SC23**) in 77% isolated yield. Oxidation of **SC23** using oxone® and aqueous NaHCO_3_ resulted in alcohol **SC33**.

#### Synthesis of C12 analogs

For the C12 derivatives, as shown in Figure 1D, mitragynine (**1**) was brominated directly in presence of NBS and AcOH to afford mainly 12 bromo mitragynine (**7**) in 47% yield. Synthesis of 12-phenyl mitragynine (**SC72**) was achieved in 67% yield using palladium-catalyzed coupling reaction of bromide **7** with phenylboronic acid. Oxone® and aqueous NaHCO_3_ mediated hydroxylation of **SC72** furnished the corresponding 7OH derivative **SC82** in 41% yield. The synthesis of 12-3’furanyl mitragynine (**SC71)** was done by a similar palladium-catalyzed reaction of bromide **7** with 3-furanylboronic acid. **SC71** on treatment with oxone® and aqueous NaHCO_3_ furnished alcohol **SC81**. The methyl group at C12 position was installed by coupling reaction of bromide **7** with DABAL-Me_3_ in presence of XPhos to afford 12-methyl mitragynine (**SC73**) in 87% isolated yield. Oxidation of **SC73** using oxone® and aqueous NaHCO_3_ yielded alcohol **SC83** in 55% yield.

#### Synthesis of individual embodiments

*methyl(E)-2-((2S,3S,12bS)-3-ethyl-8-(((trifluoromethyl)sulfonyl)oxy)-1,2,3,4,6,7,12,12b- octahydroindolo[2,3-a]quinolizin-2-yl)-3-methoxyacrylate***(4)**:*N*-Phenyl-bis(trifluoromethane sulfonimide) (66.4 mg, 0.18 mmol) was added to a solution of 9-hydroxymitragynine (65 mg, 0.16 mmol) dissolved in DCM (3 mL) under argon at RT. Et_3_N (0.07 mL, 0.50 mmol) was added to the mixture and the reaction was continued overnight. MS indicated the completion of the reaction. Then, the solvent was evaporated and the reaction mixture was diluted in EtOAc (20 mL) and was washed with brine (5x20 mL), dried over anhydrous Na_2_SO_4_ and filtered. The solvent was removed and residue was purified by flash column chromatography using 20-60%EtOAc in hexane to get the desired triflate **4** as a white solid 53 mg; (Yield, 61%). Since this is an intermediate compound, we recorded only proton NMR. ^1^H NMR (400 MHz, Chloroform-*d*) δ 8.01 (s, 1H), 7.44 (d, *J* = 0.8 Hz, 1H), 7.28 (dt, *J* = 8.0, 0.8 Hz, 1H), 7.07 (dd, *J* = 8.5, 7.6 Hz, 1H), 6.98 (d, *J* = 7.9 Hz, 1H), 3.74 (s, 3H), 3.71 (s, 3H), 3.23 – 3.10 (m, 2H), 3.02 (tt, *J* = 17.0, 4.9 Hz, 3H), 2.91 (dd, *J* = 15.4, 3.6 Hz, 1H), 2.62 – 2.51 (m, 2H), 2.47 (dd, *J* = 11.7, 3.1 Hz, 1H), 1.85 – 1.69 (m, 2H), 1.69 – 1.60 (m, 1H), 1.25 – 1.16 (m, 1H), 0.87 (t, *J* = 7.3 Hz, 3H). HRMS (ESI-TOF) *m*/*z*: [M+H]^+^ Calcd for C_23_H_28_F_3_N_2_O_6_S 517.1620; found 517.1611.

*methyl(E)-2-((2S,3S,12bS)-11-bromo-3-ethyl-8-methoxy-1,2,3,4,6,7,12,12b-octahydroindolo[2,3- a]quinolizin-2-yl)-3-methoxyacrylate* **(7)**: Mitragynine (800 mg, 2.007 mmol) was dissolved in glacial acetic acid (8 mL). Then, NBS (535.78 mg, 3.01 mmol, 1.5 eq) was added to the mixture under Argon. The mixture was stirred for 4h at RT. MS indicated the formation of bromomitragynine. The reaction mixture was basified with sat. aq. NaHCO_3_ solution and the product was extracted with DCM (3x 20 mL). The DCM layer was washed with brine (15 mL), dried over anhydrous Na_2_SO_4_ and filtered. The solvent was removed under reduced pressure and the crude product was purified by flash column chromatography using 10-25% EtOAc in hexanes. Fraction 3-18 gave 450 mg (47%) of 12-bromomitragynine (**7**) while Fr 23-38 contained 10- bromomitragynine (∼5%; with minor impurities). Since compound **7** is an intermediate compound, we recorded only proton NMR. _1_H NMR (400 MHz, Chloroform-*d*) δ 7.84 – 7.73 (m, 1H), 7.44 (s, 1H), 7.10 (d, *J* = 8.3 Hz, 1H), 6.36 (d, *J* = 8.3 Hz, 1H), 3.85 (s, 3H), 3.75 (s, 3H), 3.71 (s, 3H), 3.17 (d, *J* = 11.6 Hz, 1H), 3.12 – 2.99 (m, 3H), 2.97 – 2.87 (m, 2H), 2.57 – 2.42 (m, 3H), 1.85 (dt, *J* = 12.8, 3.1 Hz, 1H), 1.81 – 1.73 (m, 1H), 1.66 (d, *J* = 19.9 Hz, 1H), 1.25 – 1.19 (m, 1H), 0.87 (t, *J* = 7.3 Hz, 3H). HRMS (ESI-TOF) *m*/*z*: [M+H]^+^ Calcd for C_23_H_30_BrN_2_O_6_ 477.1383; found 477.1380.

*methyl(E)-2-((2S,3S,12bS)-3-ethyl-8-(furan-3-yl)-1,2,3,4,6,7,12,12b-octahydroindolo[2,3- a]quinolizin-2-yl)-3-methoxyacrylate* **(SC1)**: **4** (77.5 mg, 0.15 mmol) was dissolved in dry toluene (0.5 mL) and the solvent was removed under reduced pressure to ensure azeotropic removal of water residues. Dry methanol (1 mL) and dry toluene (1.5 mL) were added. To the resulting solution were added 3-furanylboronic acid (17.9 mg, 0.16 mmol, 1.1 equiv), K_2_CO_3_ (41.5 mg, 2 equiv) and Pd(PPh_3_)_4_ (8.7 mg, 0.05 equiv). The mixture was stirred at 80 °C for 8 hrs. The solvent was evaporated under reduced pressure and the residue was extracted with DCM (3×20 mL). The combined extracts were washed with brine (3×1/3 vol.), dried (Na_2_SO_4_) and concentrated to provide the crude product. The crude product was purified by flash column chromatography (gradient: 25-70% EtOAc in hexanes) to yield 47.6 mg (73%) of **SC1** as a yellow amorphous solid.

^1^H NMR (400 MHz, Chloroform-*d*) δ 7.81 (s, 1H), 7.55 – 7.50 (m, 1H), 7.47 (d, *J* = 1.0 Hz, 1H), 7.44 (d, *J* = 0.9 Hz, 1H), 7.30 – 7.27 (m, 1H), 7.11 (dd, *J* = 8.1, 7.2 Hz, 1H), 7.00 – 6.96 (m, 1H), 6.62 – 6.58 (m, 1H), 3.74 (s, 3H), 3.71 (s, 3H), 3.17 (d, *J* = 11.3 Hz, 1H), 3.02 (ddd, *J* = 20.5, 11.0, 2.9 Hz, 2H), 2.91 – 2.80 (m, 2H), 2.55 (q, *J* = 12.5 Hz, 1H), 2.46 – 2.36 (m, 3H), 1.84 (d, *J* = 13.4 Hz, 1H), 1.78 (dt, *J* = 13.3, 5.8 Hz, 1H), 1.63 (d, *J* = 11.4 Hz, 1H), 1.25 – 1.19 (m, 1H), 0.86 (t, *J* = 7.3 Hz, 3H). ^13^C NMR (125 MHz, Chloroform-*d*) δ 169.40, 160.75, 142.12, 140.18, 136.49, 136.30, 125.82, 125.76, 125.38, 121.33, 121.17, 113.22, 111.70, 110.25, 108.36, 61.77, 61.72, 58.00, 53.94, 51.57, 40.84, 40.17, 30.21, 24.92, 19.36, 13.03. HRMS (ESI-TOF) *m*/*z*: [M+H]^+^Calcd for C_26_H_31_N_2_O_4_ 435.2278; found 435.2273.

*methyl(E)-2-((2S,3S,12bS)-3-ethyl-8-phenyl-1,2,3,4,6,7,12,12b-octahydroindolo[2,3- a]quinolizin-2-yl)-3-methoxyacrylate* **(SC 3)**: **4** (77.5 mg, 0.15 mmol) was dissolved in dry toluene (0.5 mL) and the solvent was removed under reduced pressure to ensure azeotropic removal of water residues. Dry methanol (1 mL) and dry toluene (1.5 mL) were added. To the resulting solution were added phenylboronic acid (19.5 mg, 0.16 mmol, 1.1 equiv), K_2_CO_3_ (41.5 mg, 2 equiv) and Pd(PPh_3_)_4_ (8.7 mg, 0.05 equiv). The mixture was stirred at 80 °C for 8 hrs. The solvent was evaporated under reduced pressure and the residue was extracted with DCM (3×20 mL). The combined extracts were washed with brine (3×1/3 vol.), dried (Na_2_SO_4_) and concentrated to provide the crude product. The crude product was purified by flash column chromatography (gradient: 25-70% EtOAc in hexanes) to yield 43.3 mg (65%) of **SC3** as a light yellow amorphous solid. ^1^H NMR (400 MHz, Chloroform-*d*) δ 7.83 (s, 1H), 7.50 – 7.33 (m, 6H), 7.30 (d, *J* = 8.0 Hz, 1H), 7.15 (t, *J* = 7.7 Hz, 1H), 6.98 (d, *J* = 7.2 Hz, 1H), 3.74 (s, 3H), 3.71 (s, 3H), 3.17 (d, *J* = 11.3 Hz, 1H), 3.04 (dt, *J* = 13.1, 3.7 Hz, 1H), 2.97 (dd, *J* = 11.7, 2.2 Hz, 1H), 2.78 – 2.73 (m, 1H), 2.71 – 2.64 (m, 1H), 2.55 (q, *J* = 12.6 Hz, 1H), 2.41 (dd, *J* = 11.3, 3.1 Hz, 1H), 2.33 (td, *J* = 10.7, 10.2, 4.1 Hz, 1H), 2.01 – 1.94 (m, 1H), 1.85 (d, *J* = 12.9 Hz, 1H), 1.81 – 1.71 (m, 2H), 1.22 (d, *J* = 8.0 Hz, 1H), 0.85 (t, *J* = 7.3 Hz, 3H). ^13^C NMR (100 MHz, Chloroform-*d*) δ 169.18, 160.54, 141.67, 136.19, 136.06, 134.96, 129.77, 127.44, 126.59, 125.24, 121.00, 120.86, 111.46, 109.76, 108.17, 61.58, 61.52, 57.74, 53.69, 51.37, 40.60, 39.93, 30.00, 24.68, 19.15, 12.83. HRMS (ESI-TOF) *m*/*z*: [M+H]^+^ Calcd for C_28_H_33_N_2_O_3_ 445.2486; found 445.2484.

*methyl(E)-2-((2S,3S,12bS)-3-ethyl-8-methyl-1,2,3,4,6,7,12,12b-octahydroindolo[2,3- a]quinolizin-2-yl)-3-methoxyacrylate* **(SC4)**: Starting material **4** (77.5 mg, 0.15 mmol), Pd_2_(dba)_3_ (13.7 mg, 0.1 equiv), Xphos (10.7 mg, 0.15 equiv) and DABAL- Me3 (153.8 mg, 4 equiv) were balanced into an oven dried vial. Vial was purged with argon and dry THF (3 mL) was added under argon. Vial was sealed with a Teflon lined screw cap and heated to 60 °C. After stirring for 8 h complete conversion was observed by LC-MS. The reaction mixture was cooled to RT and concentrated in vacuo. The product was purified by flash column chromatography (gradient: 25- 75% EtOAc in hexanes); to yield 39 mg (68%) of **SC4** as a yellow solid. ^1^H NMR (400 MHz, Chloroform-*d*) δ 7.73 (s, 1H), 7.43 (s, 1H), 7.10 (d, *J* = 8.1 Hz, 1H), 6.97 (t, *J* = 7.6 Hz, 1H), 6.78 (d, *J* = 6.6 Hz, 1H), 3.72 (s, 3H), 3.71 (s, 3H), 3.29 – 3.14 (m, 2H), 3.08 – 2.93 (m, 4H), 2.62 (s, 3H), 2.59 – 2.43 (m, 3H), 1.84 – 1.74 (m, 2H), 1.64 (dd, *J* = 8.7, 5.4 Hz, 1H), 1.27 – 1.17 (m, 1H), 0.87 (t, *J* = 7.3 Hz, 3H). ^13^C NMR (100 MHz, Chloroform-*d*) δ 169.22, 160.57, 135.85, 134.92, 130.35, 126.50, 121.23, 120.38, 111.43, 108.44, 108.30, 61.55, 61.34, 57.70, 53.76, 51.37, 40.60, 39.85, 29.89, 24.43, 19.56, 19.10, 12.85. HRMS (ESI-TOF) *m*/*z*: [M+H]^+^ Calcd for C_23_H_31_N_2_O_3_ 383.2329; found 383.2327.

*methyl(E)-2-((2S,3S,7aS,12bS)-3-ethyl-7a-hydroxy-8-phenyl-1,2,3,4,6,7,7a,12b- octahydroindolo[2,3-a]quinolizin-2-yl)-3-methoxyacrylate* **(SC11)**: A saturated aq. NaHCO_3_ (3 mL) was added to a solution of **SC3** (44.4 mg, 0.10 mmol) in acetone (4 mL) at 0 °C resulting in suspension formation. A solution of oxone® (30.6 mg, 0.20 mmol) in distilled water (1 mL) was added dropwise over 5 min period (this is crucial for the reaction! slower addition is better). The reaction mixture was stirred for additional 30 min at 0 °C. Then, the content was diluted with water (2-3 mL) and the product was extracted in ethyl acetate (3x10 mL). EtOAc layer was washed with brine (15 mL), dried over anhydrous Na_2_SO_4,_ and filtered. Solvent was removed under reduced pressure and the content was purified by flash column chromatography (gradient: 25-65% EtOAc in hexanes); to yield 15.2 mg (33%) of **SC11** as a white solid. _1_H NMR (500 MHz, Chloroform- *d*) δ 7.59 – 7.51 (m, 3H), 7.46 – 7.41 (m, 3H), 7.41 – 7.35 (m, 2H), 7.13 (dd, *J* = 7.8, 0.9 Hz, 1H), 3.81 (s, 3H), 3.69 (s, 3H), 3.09 (dd, *J* = 11.1, 2.6 Hz, 1H), 3.04 – 2.95 (m, 2H), 2.81 (td, *J* = 13.6, 11.0 Hz, 1H), 2.64 – 2.56 (m, 1H), 2.46 (ddd, *J* = 11.9, 4.7, 2.3 Hz, 1H), 2.41 (dd, *J* = 11.4, 3.0 Hz, 1H), 2.05 (d, *J* = 1.5 Hz, 1H), 1.94 (ddt, *J* = 30.4, 13.8, 2.7 Hz, 2H), 1.67 (ddd, *J* = 13.5, 11.2, 7.0 Hz, 1H), 1.57 (d, *J* = 11.2 Hz, 1H), 1.49 (td, *J* = 13.3, 4.1 Hz, 1H), 1.28 – 1.19 (m, 1H), 0.79 (t, *J* = 7.3 Hz, 3H). ^13^C NMR (100 MHz, Chloroform-*d*) δ 183.99, 169.30, 160.76, 154.17, 139.44, 139.33, 137.29, 129.61, 129.33, 128.11, 127.61, 127.57, 120.36, 111.24, 80.98, 61.80, 61.48, 58.15, 51.32, 50.07, 40.47, 39.21, 34.85, 26.07, 18.93, 12.78. HRMS (ESI-TOF) *m*/*z*: [M+H]^+^Calcd for C_28_H_33_N_2_O4 461.2435; found 461.2431.

*methyl(E)-2-((2S,3S,7aS,12bS)-3-ethyl-7a-hydroxy-8-methyl-1,2,3,4,6,7,7a,12b- octahydroindolo[2,3-a]quinolizin-2-yl)-3-methoxyacrylate* **(SC12)**: A saturated aq. NaHCO_3_ (3 mL) was added to a solution of **SC4** (38.2 mg, 0.10 mmol) in acetone (4 mL) at 0 °C resulting in suspension formation. A solution of oxone® (30.6 mg, 0.20 mmol) in distilled water (1 mL) was added dropwise over 5 min period. The reaction mixture was stirred for additional 30 min at 0 °C. Then, the content was diluted with water (2-3 mL) and the product was extracted in ethyl acetate (3x10 mL). EtOAc layer was washed with brine (15 mL), dried over anhydrous Na_2_SO_4,_ and filtered. Solvent was removed under reduced pressure and the content was purified by flash column chromatography (gradient: 25-65% EtOAc in hexanes); to yield 16.7 mg (42%) of **SC12** as a white solid. ^1^H NMR (400 MHz, Chloroform-*d*) δ 7.44 (s, 1H), 7.38 (d, *J* = 7.6 Hz, 1H), 7.21 (t, *J* = 7.6 Hz, 1H), 6.95 (d, *J* = 7.7 Hz, 1H), 3.81 (s, 3H), 3.67 (s, 3H), 3.13 (dd, *J* = 10.7, 2.4 Hz, 1H), 3.08 – 2.97 (m, 2H), 2.80 (ddd, *J* = 21.6, 12.8, 10.0 Hz, 2H), 2.68 – 2.59 (m, 2H), 2.48 (d, *J* = 11.6 Hz, 1H), 2.45 (s, 3H), 2.09 – 2.07(m, 1H), 1.86 (d, *J* = 13.2 Hz, 1H), 1.68 – 1.55 (m, 3H), 1.21 (dd, *J* = 7.2, 5.6 Hz, 1H), 0.82 (t, *J* = 7.3 Hz, 3H). ^13^C NMR (100 MHz, Chloroform-*d*) δ 184.01, 169.32, 160.78, 153.58, 138.01, 134.44, 129.39, 127.89, 118.78, 111.20, 81.30, 61.81, 61.44, 58.16, 51.30, 50.11, 40.51, 39.32, 34.94, 26.06, 18.95, 17.11, 12.82. HRMS (ESI-TOF) *m*/*z*: [M+H]^+^ Calcd for C_23_H_31_N_2_O_4_ 399.227; found 399.2277.

*methyl(E)-2-((2S,3S,7aS,12bS)-3-ethyl-8-(furan-3-yl)-7a-hydroxy-1,2,3,4,6,7,7a,12b- octahydroindolo[2,3-a]quinolizin-2-yl)-3-methoxyacrylate* **(SC13)**: A saturated aq. NaHCO_3_ (3 mL) was added to a solution of **SC1** (43.4 mg, 0.10 mmol) in acetone (4 mL) at 0 °C resulting in suspension formation. A solution of oxone® (30.6 mg, 0.20 mmol) in distilled water (1 mL) was added dropwise over 5 min period. The reaction mixture was stirred for additional 30 min at 0 °C. Then, the content was diluted with water (2-3 mL) and the product was extracted in ethyl acetate (3x10 mL). EtOAc layer was washed with brine (15 mL), dried over anhydrous Na_2_SO_4,_ and filtered. Solvent was removed under reduced pressure and the content was purified by flash column chromatography (gradient: 25-65% EtOAc in hexanes); to yield 17.1 mg (38%) of **SC13** as a white solid. ^1^H NMR (500 MHz, Chloroform-*d*) δ 8.05 (dd, *J* = 1.6, 0.9 Hz, 1H), 7.50 – 7.48 (m, 2H), 7.44 (s, 1H), 7.34 (t, *J* = 7.7 Hz, 1H), 7.24 (dd, *J* = 7.8, 1.0 Hz, 1H), 6.87 (dd, *J* = 1.9, 0.9 Hz, 1H), 3.82 (s, 3H), 3.68 (s, 3H), 3.14 (dd, *J* = 11.0, 2.5 Hz, 1H), 3.06 – 2.99 (m, 2H), 2.88 – 2.71 (m, 2H), 2.62 – 2.55 (m, 2H), 2.46 (dd, *J* = 11.4, 3.1 Hz, 1H), 2.22 (s, 1H), 1.90 (dd, *J* = 13.6, 3.2 Hz, 1H), 1.68 (ddt, *J* = 14.1, 11.8, 7.1 Hz, 2H), 1.45 (td, *J* = 13.5, 12.8, 4.7 Hz, 1H), 1.26 – 1.19 (m, 1H), 0.81 (t, *J* = 7.3 Hz, 3H). ^13^C NMR (100 MHz, Chloroform-*d*) δ 184.10, 169.31, 160.80, 154.40, 142.99, 141.72, 136.55, 129.85, 129.63, 126.11, 123.43, 120.12, 111.19, 111.12, 81.19, 61.81, 61.40, 58.14, 51.31, 50.05, 40.48, 39.30, 32.67, 26.09, 18.95, 12.81. HRMS (ESI-TOF) *m*/*z*: [M+H]^+^ Calcd for C_26_H_31_N_2_O_5_ 451.2227; found 451.2224.

*methyl(E)-2-((2S,3S,12bS)-3-ethyl-8-methoxy-9-phenyl-1,2,3,4,6,7,12,12b-octahydroindolo[2,3- a]quinolizin-2-yl)-3-methoxyacrylate* **(SC21)**: Starting material **6** (71.6 mg, 0.15 mmol), phenylboronic acid (40.2 mg, 2.2 equiv), KOAc (33.8 mg, 2.3 equiv) and Pd(dppf)Cl_2_·CH_2_Cl_2_ (6.1 mg, 0.05 equiv) were balanced into an oven-dried vial. Vial was purged with argon and dry THF (3 mL) was added under a stream of argon. Vial was closed with a Teflon lined solid screw cap and heated to 70 °C. After 6h LC-MS and TLC indicated full consumption of starting material. The solvent was evaporated under reduced pressure and the residue was extracted with DCM (3×20 mL). The combined extracts were washed with brine (3×1/3 vol.), dried (Na_2_SO_4_) and concentrated to provide the crude product. The crude product was purified by flash column chromatography (gradient: 25-70% EtOAc in hexanes) to yield 50.5 mg (71%) of **SC21** as a yellow amorphous solid. ^1^H NMR (400 MHz, Chloroform-*d*) δ 7.77 (s, 1H), 7.65 – 7.58 (m, 2H), 7.41 (dd, *J* = 15.3, 7.8 Hz, 3H), 7.30 (d, *J* = 7.5 Hz, 1H), 7.14 – 7.06 (m, 2H), 3.74 (s, 3H), 3.72 (s, 3H), 3.51 (s, 3H), 3.19 (d, *J* = 11.9 Hz, 2H), 3.09 – 2.93 (m, 4H), 2.63 – 2.45 (m, 3H), 1.87 – 1.75 (m, 2H), 1.64 (d, *J* = 11.4 Hz, 1H), 1.25 – 1.17 (m, 1H), 0.88 (t, *J* = 7.3 Hz, 3H). ^13^C NMR (100 MHz, Chloroform-*d*) δ 169.18, 160.53, 151.01, 139.65, 137.39, 135.53, 129.55, 128.05, 126.13, 125.13, 124.33, 121.50, 111.49, 107.48, 107.06, 61.65, 61.57, 61.32, 57.81, 53.75, 51.37, 40.67, 39.92, 29.92, 23.57, 19.14, 12.88. HRMS (ESI-TOF) *m*/*z*: [M+H]^+^ Calcd for C_29_H_35_N_2_O4 475.2591; found 475.2586.

*methyl(E)-2-((2S,3S,12bS)-3-ethyl-9-(furan-3-yl)-8-methoxy-1,2,3,4,6,7,12,12b- octahydroindolo[2,3-a]quinolizin-2-yl)-3-methoxyacrylate* **(SC22)**: Starting material **6** (71.6 mg, 0.15 mmol), 3-furanylboronic acid (36.9 mg, 2.2 equiv), KOAc (33.8 mg, 2.3 equiv) and Pd(dppf)Cl_2_·CH_2_Cl_2_ (6.1 mg, 0.05 equiv) were balanced into an oven-dried vial. Vial was purged with argon and dry THF (3 mL) was added under a stream of argon. Vial was closed with a Teflon lined solid screw cap and heated to 70 °C. After 6h LC-MS and TLC indicated full consumption of starting material. The solvent was evaporated under reduced pressure and the residue was extracted with DCM (3×20 mL). The combined extracts were washed with brine (3×1/3 vol.), dried (Na_2_SO4) and concentrated to provide the crude product. The crude product was purified by flash column chromatography (gradient: 25-70% EtOAc in hexanes) to yield 41.1 mg (59%) of **SC22** as a light yellow solid. ^1^H NMR (400 MHz, Chloroform-*d*) δ 7.76 (s, 1H), 7.68 (s, 1H), 7.56 – 7.51 (m, 1H), 7.43 (s, 1H), 7.03 (d, *J* = 7.9 Hz, 1H), 6.71 – 6.67 (m, 1H), 6.50 (d, *J* = 8.4 Hz, 1H), 3.89 (s, 3H), 3.73 (s, 3H), 3.70 (s, 3H), 3.19 – 2.90 (m, 6H), 2.58 – 2.43 (m, 3H), 1.82 – 1.74 (m, 2H), 1.63 (br s, 1H), 1.23 – 1.16 (m, 1H), 0.87 (t, *J* = 7.3 Hz, 3H). ^13^C NMR (100 MHz, Chloroform-*d*) δ 169.18, 160.56, 154.05, 143.49, 138.21, 134.97, 133.97, 123.82, 121.18, 117.80, 111.44, 110.33, 109.48, 108.79, 100.22, 61.57, 61.36, 57.79, 55.43, 53.70, 51.35, 40.75, 39.84, 29.97, 23.87, 19.15, 12.88.. HRMS (ESI-TOF) *m*/*z*: [M+H]^+^ Calcd for C_27_H_33_N_2_O_5_ 465.2384; found 465.2381.

*methyl(E)-2-((2S,3S,12bS)-3-ethyl-8-methoxy-9-methyl-1,2,3,4,6,7,12,12b-octahydroindolo[2,3- a]quinolizin-2-yl)-3-methoxyacrylate* **(SC23)**: Starting material **6** (71.6 mg, 0.15 mmol), Pd_2_(dba)_3_ (13.7 mg, 0.1 equiv), Xphos (10.7 mg, 0.15 equiv) and DABAL- Me3 (153.8 mg, 4 equiv) were balanced into an oven dried vial. Vial was purged with argon and dry THF (3 mL) was added under argon. Vial was sealed with a Teflon lined screw cap and heated to 60 °C. After stirring for 8 h complete conversion was observed by LC-MS. The reaction mixture was cooled to RT and concentrated in vacuo. The product was purified by flash column chromatography (gradient: 25- 75% EtOAc in hexanes); to yield 31.8 mg (77%) of **SC23** as a yellow solid. ^1^H NMR (400 MHz, Chloroform-*d*) δ 7.68 (s, 1H), 7.43 (s, 1H), 6.97 (d, *J* = 10.3 Hz, 1H), 6.88 (d, *J* = 8.1 Hz, 1H), 3.84 (s, 3H), 3.73 (s, 3H), 3.71 (s, 3H), 3.16 (d, *J* = 10.8 Hz, 2H), 3.08 – 3.00 (m, 2H), 2.95 (d, *J* = 12.5 Hz, 2H), 2.58 – 2.43 (m, 3H), 2.34 (s, 3H), 1.82 – 1.75 (m, 2H), 1.63 (d, *J* = 11.8 Hz, 1H), 1.23 – 1,19 (m, 1H), 0.88 (t, *J* = 7.2 Hz, 3H). ^13^C NMR (100 MHz, Chloroform-*d*) δ 169.21, 160.54, 151.43, 136.58, 135.03, 124.29, 121.29, 119.95, 111.46, 106.80, 106.47, 61.74, 61.57, 61.40, 57.82, 53.80, 51.36, 40.64, 39.94, 29.85, 23.46, 19.14, 15.10, 12.88. HRMS (ESI-TOF) *m*/*z*: [M+H]^+^ Calcd for C_24_H_33_N_2_O_4_ 413.2435; found 413.2433.

*methyl(E)-2-((2S,3S,7aS,12bS)-3-ethyl-7a-hydroxy-8-methoxy-9-phenyl-1,2,3,4,6,7,7a,12b- octahydroindolo[2,3-a]quinolizin-2-yl)-3-methoxyacrylate* **(SC31)**: A saturated aq. NaHCO_3_ (3 mL) was added to a solution of **SC21** (47.4 mg, 0.10 mmol) in acetone (4 mL) at 0 °C resulting in suspension formation. A solution of oxone® (30.6 mg, 0.20 mmol) in distilled water (1 mL) was added dropwise over 5 min period. The reaction mixture was stirred for additional 30 min at 0 °C. Then, the content was diluted with water (2-3 mL) and the product was extracted in ethyl acetate (3x10 mL). EtOAc layer was washed with brine (15 mL), dried over anhydrous Na_2_SO_4,_ and filtered. Solvent was removed under reduced pressure and the content was purified by flash column chromatography (gradient: 25-65% EtOAc in hexanes); to yield 15.1 mg (31%) of **SC31** as a white solid. ^1^H NMR (400 MHz, Chloroform-*d*) δ 7.60 – 7.54 (m, 2H), 7.47 – 7.30 (m, 6H), 3.82 (s, 3H), 3.70 (s, 3H), 3.47 (s, 3H), 3.18 – 3.12 (m, 1H), 3.05 (t, *J* = 13.0 Hz, 2H), 2.89 – 2.78 (m, 2H), 2.66 (t, *J* = 14.2 Hz, 2H), 2.53 – 2.47 (m, 1H), 2.30 (s, 1H), 1.91 (d, *J* = 13.6 Hz, 1H), 1.86 – 1.67 (m, 3H), 1.25 (d, *J* = 5.5 Hz, 1H), 0.83 (t, *J* = 7.3 Hz, 3H). ^13^C NMR (100 MHz, Chloroform-*d*) δ 184.21, 169.30, 160.75, 154.66, 154.25, 138.06, 132.92, 132.81, 132.36, 128.93, 128.42, 127.23, 117.27, 111.28, 81.10, 61.81, 61.60, 61.43, 58.21, 51.31, 50.13, 40.55, 39.28, 36.35, 26.10, 18.97, 12.83. HRMS (ESI-TOF) *m*/*z*: [M+H]^+^ Calcd for C_29_H_35_N_2_O_5_ 491.2540; found 491.2542.

*methyl(E)-2-((2S,3S,7aS,12bS)-3-ethyl-9-(furan-3-yl)-7a-hydroxy-8-methoxy-1,2,3,4,6,7,7a,12b- octahydroindolo[2,3-a]quinolizin-2-yl)-3-methoxyacrylate* **(SC32)**: A saturated aq. NaHCO_3_ (3 mL) was added to a solution of **SC22** (46.4 mg, 0.10 mmol) in acetone (4 mL) at 0 °C resulting in suspension formation. A solution of oxone® (30.6 mg, 0.20 mmol) in distilled water (1 mL) was added dropwise over 5 min period. The reaction mixture was stirred for additional 30 min at 0 °C. Then, the content was diluted with water (2-3 mL) and the product was extracted in ethyl acetate (3x10 mL). EtOAc layer was washed with brine (15 mL), dried over anhydrous Na_2_SO_4,_ and filtered. Solvent was removed under reduced pressure and the content was purified by flash column chromatography (gradient: 25-65% EtOAc in hexanes); to yield 16.8 mg (35%) of **SC32** as a white solid. ^1^H NMR (400 MHz, Chloroform-*d*) δ 8.36 (s, 1H), 7.50 – 7.38 (m, 3H), 6.92 (s, 1H), 6.76 (d, *J* = 8.5 Hz, 1H), 3.89 (s, 3H), 3.84 (s, 3H), 3.72 (s, 3H), 3.11 (d, *J* = 10.7 Hz, 1H), 3.07 – 2.98 (m, 2H), 2.96 – 2.89 (m, 1H), 2.80 (t, *J* = 12.1 Hz, 1H), 2.64 (d, *J* = 12.2 Hz, 2H), 2.49 (d, *J* = 11.4 Hz, 1H), 1.95 (d, *J* = 13.6 Hz, 1H), 1.76 – 1.61 (m, 4H), 1.21 (t, *J* = 6.7 Hz, 1H), 0.82 (t, *J* = 7.2 Hz, 3H). ^13^C NMR (100 MHz, Chloroform-*d*) δ 183.61, 169.56, 160.82, 154.70, 151.65, 142.60, 141.92, 128.39, 127.51, 122.41, 118.88, 111.85, 109.50, 109.29, 81.28, 61.69, 61.64, 58.19, 55.79, 51.49, 50.21, 40.62, 39.71, 36.54, 26.49, 19.17, 13.05. HRMS (ESI-TOF) *m*/*z*: [M+H]^+^ Calcd for C_27_H_33_N_2_O_6_ 481.2333; found 481.2328.

*methyl(E)-2-((2S,3S,7aS,12bS)-3-ethyl-7a-hydroxy-8-methoxy-9-methyl-1,2,3,4,6,7,7a,12b- octahydroindolo[2,3-a]quinolizin-2-yl)-3-methoxyacrylate* **(SC33)**: A saturated aq. NaHCO_3_ (3 mL) was added to a solution of **SC23** (41.2 mg, 0.10 mmol) in acetone (4 mL) at 0 °C resulting in suspension formation. A solution of oxone® (30.6 mg, 0.20 mmol) in distilled water (1 mL) was added dropwise over 5 min period. The reaction mixture was stirred for additional 30 min at 0 °C. Then, the content was diluted with water (2-3 mL) and the product was extracted in ethyl acetate (3x10 mL). EtOAc layer was washed with brine (15 mL), dried over anhydrous Na_2_SO_4,_ and filtered. Solvent was removed under reduced pressure and the content was purified by flash column chromatography (gradient: 25-65% EtOAc in hexanes); to yield 18.8 mg (44%) of **SC33** as a white solid. ^1^H NMR (400 MHz, Chloroform-*d*) δ 7.44 (s, 1H), 7.23 (d, *J* = 7.7 Hz, 1H), 7.14 (d, *J* = 7.7 Hz, 1H), 3.89 (s, 3H), 3.81 (s, 3H), 3.68 (s, 3H), 3.14 – 2.96 (m, 3H), 2.86 – 2.74 (m, 2H), 2.70 –2.59 (m, 2H), 2.47 (dd, *J* = 11.5, 3.1 Hz, 1H), 2.29 (s, 3H), 1.87 (d, *J* = 13.6 Hz, 1H), 1.77 – 1.64 (m, 3H), 1.59 (d, *J* = 11.7 Hz, 1H), 1.24 – 1.18 (m, 1H), 0.82 (t, *J* = 7.3 Hz, 3H). ^13^C NMR (100 MHz, Chloroform-*d*) δ 183.05, 169.32, 160.76, 155.27, 153.54, 132.35, 131.68, 129.29, 116.99, 111.24, 80.92, 61.79, 61.73, 61.50, 58.19, 51.30, 50.19, 40.52, 39.30, 36.06, 26.10, 18.95, 15.77, 12.81. HRMS (ESI-TOF) *m*/*z*: [M+H]^+^ Calcd for C_24_H_32_N_2_O_5_ 429.2384; found 429.2380.

*methyl(E)-2-((2S,3S,12bS)-3-ethyl-11-(furan-3-yl)-8-methoxy-1,2,3,4,6,7,12,12b- octahydroindolo[2,3-a]quinolizin-2-yl)-3-methoxyacrylate* **(SC71)**: **7** (71.6 mg, 0.15 mmol) was dissolved in dry toluene (0.5 mL) and the solvent was removed under reduced pressure to ensure azeotropic removal of water residues. Dry methanol (1 mL) and dry toluene (1.5 mL) were added. To the resulting solution were added 3-furanylboronic acid (17.9 mg, 0.16 mmol, 1.1 equiv), K_2_CO_3_ (41.5 mg, 2 equiv) and Pd(PPh_3_)_4_ (8.7 mg, 0.05 equiv). The mixture was stirred at 80 °C for 8 hrs. The solvent was evaporated under reduced pressure and the residue was extracted with DCM (3×20 mL). The combined extracts were washed with brine (3×1/3 vol.), dried (Na_2_SO_4_) and concentrated to provide the crude product. The crude product was purified by flash column chromatography (gradient: 25-70% EtOAc in hexanes) to yield 48 mg (69 %) of **SC71** as a yellow amorphous solid. ^1^H NMR (500 MHz, Chloroform-*d*) δ 7.75 (s, 1H), 7.68 (dd, *J* = 1.5, 0.9 Hz, 1H), 7.54 (t, *J* = 1.7 Hz, 1H), 7.43 (s, 1H), 7.03 (d, *J* = 8.0 Hz, 1H), 6.70 (dd, *J* = 1.8, 0.9 Hz, 1H), 6.50 (d, *J* = 8.0 Hz, 1H), 3.90 (s, 3H), 3.73 (s, 3H), 3.70 (s, 3H), 3.21 – 3.09 (m, 2H), 3.09 – 2.96 (m, 3H), 2.93 (dd, *J* = 11.3, 5.6 Hz, 1H), 2.58 – 2.44 (m, 3H), 1.84 – 1.74 (m, 2H), 1.62 (s, 1H), 1.21 (dddd, *J* = 13.5, 7.4, 3.3, 1.0 Hz, 1H), 0.87 (t, *J* = 7.4 Hz, 3H). ^13^C NMR (100 MHz, Chloroform-*d*) δ 169.18, 160.59, 154.07, 143.53, 138.19, 135.00, 134.01, 123.84, 121.19, 117.83, 111.48, 110.31, 109.51, 108.81, 100.22, 61.53, 61.38, 57.81, 55.48, 53.71, 51.31, 40.78, 39.87, 30.00, 23.90, 19.17, 12.89. HRMS (ESI-TOF) *m*/*z*: [M+H]^+^ Calcd for C_27_H_33_N_2_O_5_ 465.2384; found 465.2383.

*methyl(E)-2-((2S,3S,12bS)-3-ethyl-8-methoxy-11-phenyl-1,2,3,4,6,7,12,12b- octahydroindolo[2,3-a]quinolizin-2-yl)-3-methoxyacrylate* **(SC72)**: **7** (71.6 mg, 0.15 mmol) was dissolved in dry toluene (0.5 mL) and the solvent was removed under reduced pressure to ensure azeotropic removal of water residues. Dry methanol (1 mL) and dry toluene (1.5 mL) were added. To the resulting solution were added phenylboronic acid (19.5 mg, 0.16 mmol, 1.1 equiv), K_2_CO_3_ (41.5 mg, 2 equiv) and Pd(PPh_3_)_4_ (8.7 mg, 0.05 equiv). The mixture was stirred at 80 °C for 8 hrs. The solvent was evaporated under reduced pressure and the residue was extracted with DCM (3×20 mL). The combined extracts were washed with brine (3×1/3 vol.), dried (Na_2_SO_4_) and concentrated to provide the crude product. The crude product was purified by flash column chromatography (gradient: 25-70% EtOAc in hexanes) to yield 47.7 mg (67%) of **SC72** as a light yellow solid. ^1^H NMR (400 MHz, Chloroform-*d*) δ 7.87 (s, 1H), 7.59 (dd, *J* = 8.0, 1.4 Hz, 2H), 7.47 (t, *J* = 7.6 Hz, 2H), 7.40 (d, *J* = 0.8 Hz, 1H), 7.37 – 7.32 (m, 1H), 7.02 (d, *J* = 7.9 Hz, 1H), 6.55 (d, *J* = 8.0 Hz, 1H), 3.91 (s, 3H), 3.70 (s, 3H), 3.68 (s, 3H), 3.20 – 3.11 (m, 2H), 3.06 – 2.98 (m, 3H), 2.93 (dd, *J* = 11.3, 5.5 Hz, 1H), 2.59 – 2.51 (m, 1H), 2.46 (dd, *J* = 11.3, 2.7 Hz, 2H), 1.82 – 1.69 (m, 2H), 1.61 (s, 1H), 1.28 – 1.16 (m, 1H), 0.86 (t, *J* = 7.4 Hz, 3H). ^13^C NMR (100 MHz, Chloroform-*d*) δ 169.15, 160.47, 154.16, 139.68, 134.72, 133.93, 127.99, 127.99, 126.63, 122.03, 118.82, 117.75, 111.42, 108.55, 100.35, 61.47, 61.39, 57.80, 55.35, 53.70, 51.25, 40.75, 39.81, 29.93, 23.90, 19.13, 12.85. HRMS (ESI-TOF) *m*/*z*: [M+H]^+^ Calcd for C_29_H_34_N_2_O_4_ 475.2591; found 475.2590.

*methyl(E)-2-((2S,3S,12bS)-3-ethyl-8-methoxy-11-methyl-1,2,3,4,6,7,12,12b- octahydroindolo[2,3-a]quinolizin-2-yl)-3-methoxyacrylate* **(SC73)**: Starting material **7** (71.6 mg, 0.15 mmol), Pd_2_(dba)_3_ (13.7 mg, 0.1 equiv), Xphos (10.7 mg, 0.15 equiv) and DABAL- Me3 (153.8 mg, 4 equiv) were balanced into an oven dried vial. Vial was purged with argon and dry THF (3 mL) was added under argon. Vial was sealed with a Teflon lined screw cap and heated to 60 °C. After stirring for 8 h complete conversion was observed by LC-MS. The reaction mixture was cooled to RT and concentrated in vacuo. The product was purified by flash column chromatography (gradient: 25-75% EtOAc in hexanes); to yield 35.9 mg (87%) of **SC73** as a yellow solid. ^1^H NMR (500 MHz, Chloroform-*d*) δ 7.52 (s, 1H), 7.44 (s, 1H), 6.78 (dd, *J* = 7.8, 1.0 Hz, 1H), 6.38 (d, *J* = 7.8 Hz, 1H), 3.85 (s, 3H), 3.74 (s, 3H), 3.71 (s, 3H), 3.18 (dd, *J* = 11.4, 2.3 Hz, 1H), 3.15 – 3.00 (m, 3H), 2.99 – 2.88 (m, 2H), 2.57 – 2.43 (m, 3H), 2.37 (s, 3H), 1.87 – 1.75 (m, 2H), 1.62 (d, *J* = 10.6 Hz, 1H), 1.21 (dddd, *J* = 12.3, 11.2, 5.0, 2.8 Hz, 1H), 0.88 (d, *J* = 7.3 Hz, 3H). ^13^C NMR (100 MHz, Chloroform-*d*) δ 169.20, 160.48, 152.93, 136.41, 133.48, 121.86, 117.11, 112.85, 111.57, 108.50, 99.90, 61.54, 61.38, 57.79, 55.49, 53.75, 51.35, 40.74, 39.87, 30.00, 23.87, 19.11, 16.07, 12.86. HRMS (ESI-TOF) *m*/*z*: [M+H]^+^ Calcd for C_24_H_33_N_2_O_4_ 413.2435; found 413.2435.

*methyl(E)-2-((2S,3S,7aS,12bS)-3-ethyl-11-(furan-3-yl)-7a-hydroxy-8-methoxy- 1,2,3,4,6,7,7a,12b-octahydroindolo[2,3-a]quinolizin-2-yl)-3-methoxyacrylate* **(SC81)**: A saturated aq. NaHCO_3_ (3 mL) was added to a solution of **SC71** (46.4 mg, 0.10 mmol) in acetone (4 mL) at 0 °C resulting in suspension formation. A solution of oxone® (30.6 mg, 0.20 mmol) in distilled water (1 mL) was added dropwise over 5 min period. The reaction mixture was stirred for additional 30 min at 0 °C. Then, the content was diluted with water (2-3 mL) and the product was extracted in ethyl acetate (3x10 mL). EtOAc layer was washed with brine (15 mL), dried over anhydrous Na_2_SO_4,_ and filtered. Solvent was removed under reduced pressure and the content was purified by flash column chromatography (gradient: 25-65% EtOAc in hexanes); to yield 21.6 mg (45%) of **SC81** as a white solid. ^1^H NMR (500 MHz, Chloroform-*d*) δ 8.35 (dd, *J* = 1.6, 0.8 Hz, 1H), 7.46 (s, 1H), 7.45 – 7.42 (m, 2H), 6.92 (dd, *J* = 1.9, 0.8 Hz, 1H), 6.75 (d, *J* = 8.6 Hz, 1H), 3.88 (s, 3H), 3.83 (s, 3H), 3.72 (s, 3H), 3.11 (dd, *J* = 11.0, 2.6 Hz, 1H), 3.05 – 2.98 (m, 2H), 2.96 – 2.87 (m, 1H), 2.84 – 2.77 (m, 1H), 2.63 (ddt, *J* = 13.8, 6.5, 2.3 Hz, 2H), 2.49 (ddd, *J* = 11.4, 3.1, 1.0 Hz, 1H), 2.18 (s, 1H), 1.95 (dtd, *J* = 13.5, 3.0, 1.2 Hz, 1H), 1.74 – 1.63 (m, 3H), 1.25 – 1.20 (m, 1H), 0.82 (t, *J* = 7.3 Hz, 3H). ^13^C NMR (100 MHz, Chloroform-*d*) δ 183.61, 169.56, 160.82, 154.70, 151.65, 142.60, 141.92, 128.39, 127.51, 122.41, 118.88, 111.85, 109.50, 109.29, 81.28, 61.69, 61.64, 58.19, 55.79, 51.49, 50.21, 40.62, 39.71, 36.54, 26.49, 19.17, 13.05. HRMS (ESI-TOF) *m*/*z*: [M+H]^+^ Calcd for C_27_H_33_N_2_O_6_ 481.2333; found 481.2329.

*methyl(E)-2-((2S,3S,7aS,12bS)-3-ethyl-7a-hydroxy-8-methoxy-11-phenyl-1,2,3,4,6,7,7a,12b- octahydroindolo[2,3-a]quinolizin-2-yl)-3-methoxyacrylate* **(SC82)**: A saturated aq. NaHCO_3_ (3 mL) was added to a solution of **SC72** (47.4 mg, 0.10 mmol) in acetone (4 mL) at 0 °C resulting in suspension formation. A solution of oxone® (30.6 mg, 0.20 mmol) in distilled water (1 mL) was added dropwise over 5 min period. The reaction mixture was stirred for additional 30 min at 0 °C. Then, the content was diluted with water (2-3 mL) and the product was extracted in ethyl acetate (3x10 mL). EtOAc layer was washed with brine (15 mL), dried over anhydrous Na_2_SO_4,_ and filtered. Solvent was removed under reduced pressure and the content was purified by flash column chromatography (gradient: 25-65% EtOAc in hexanes); to yield 20.1 mg (41%) of **SC82** as a white solid. ^1^H NMR (400 MHz, Chloroform-*d*) δ 7.87 (d, *J* = 7.1 Hz, 2H), 7.48 (d, *J* = 8.6 Hz, 1H), 7.40 (dd, *J* = 14.8, 7.2 Hz, 3H), 7.30 (d, *J* = 7.4 Hz, 1H), 6.82 (d, *J* = 8.6 Hz, 1H), 3.91 (s, 3H), 3.78 (s, 3H), 3.70 (s, 3H), 3.13 – 3.07 (m, 1H), 3.06 – 2.94 (m, 2H), 2.83 (dt, *J* = 21.6, 12.1 Hz, 2H), 2.70 – 2.61 (m, 2H), 2.48 (d, *J* = 11.4 Hz, 1H), 1.90 (d, *J* = 13.8 Hz, 1H), 1.79 – 1.61 (m, 4H), 1.21 (d, *J* = 7.2 Hz, 1H), 0.82 (t, *J* = 7.2 Hz, 3H). ^13^C NMR (100 MHz, Chloroform-*d*) δ 183.40, 169.36, 160.56, 155.07, 151.97, 137.82, 130.84, 129.44, 127.86, 127.26, 127.12, 126.64, 111.54, 109.30, 80.99, 61.47, 61.33, 57.99, 55.51, 51.20, 50.02, 40.34, 39.47, 36.19, 26.06, 18.93, 12.82. HRMS (ESI-TOF) *m*/*z*: [M+H]^+^ Calcd for C_29_H_35_N_2_O_5_ 491.2540; found 491.2542.

*methyl(E)-2-((2S,3S,7aS,12bS)-3-ethyl-7a-hydroxy-8-methoxy-11-methyl-1,2,3,4,6,7,7a,12b- octahydroindolo[2,3-a]quinolizin-2-yl)-3-methoxyacrylate* **(SC83)**: A saturated aq. NaHCO_3_ (3 mL) was added to a solution of **SC73** (41.2 mg, 0.10 mmol) in acetone (4 mL) at 0 °C resulting in suspension formation. A solution of oxone® (30.6 mg, 0.20 mmol) in distilled water (1 mL) was added dropwise over 5 min period. The reaction mixture was stirred for additional 30 min at 0 °C. Then, the content was diluted with water (2-3 mL) and the product was extracted in ethyl acetate (3x10 mL). EtOAc layer was washed with brine (15 mL), dried over anhydrous Na_2_SO_4,_ and filtered. Solvent was removed under reduced pressure and the content was purified by flash column chromatography (gradient: 25-65% EtOAc in hexanes); to yield 23.6 mg (55%) of **SC83** as a white solid. ^1^H NMR (400 MHz, Chloroform-*d*) δ 7.44 (s, 1H), 7.04 (d, *J* = 7.6 Hz, 1H), 6.61 (d, *J* = 8.6 Hz, 1H), 3.82 (s, 3H), 3.81 (s, 3H), 3.71 (s, 3H), 3.04 (ddd, *J* = 13.6, 10.9, 7.5 Hz, 3H), 2.87 – 2.73 (m, 2H), 2.66 – 2.56 (m, 2H), 2.50 – 2.45 (m, 1H), 2.44 (s, 3H), 1.94 – 1.87 (m, 1H), 1.77 – 1.66 (m, 2H), 1.66 – 1.56 (m, 2H), 1.27 – 1.21 (m, 1H), 0.82 (t, *J* = 7.3 Hz, 3H). ^13^C NMR (100 MHz, Chloroform-*d*) δ 182.80, 169.29, 160.52, 153.91, 153.03, 131.77, 126.22, 123.41, 111.56, 108.67, 81.22, 61.66, 61.55, 58.18, 55.42, 51.27, 50.10, 40.52, 39.28, 35.85, 25.98, 18.98, 15.73, 12.81. HRMS (ESI-TOF) *m*/*z*: [M+H]^+^ Calcd for C_24_H_33_N_2_O_5_ 429.2384; found 429.2380.

## Appendix 1

**Appendix 1-Table 1.**
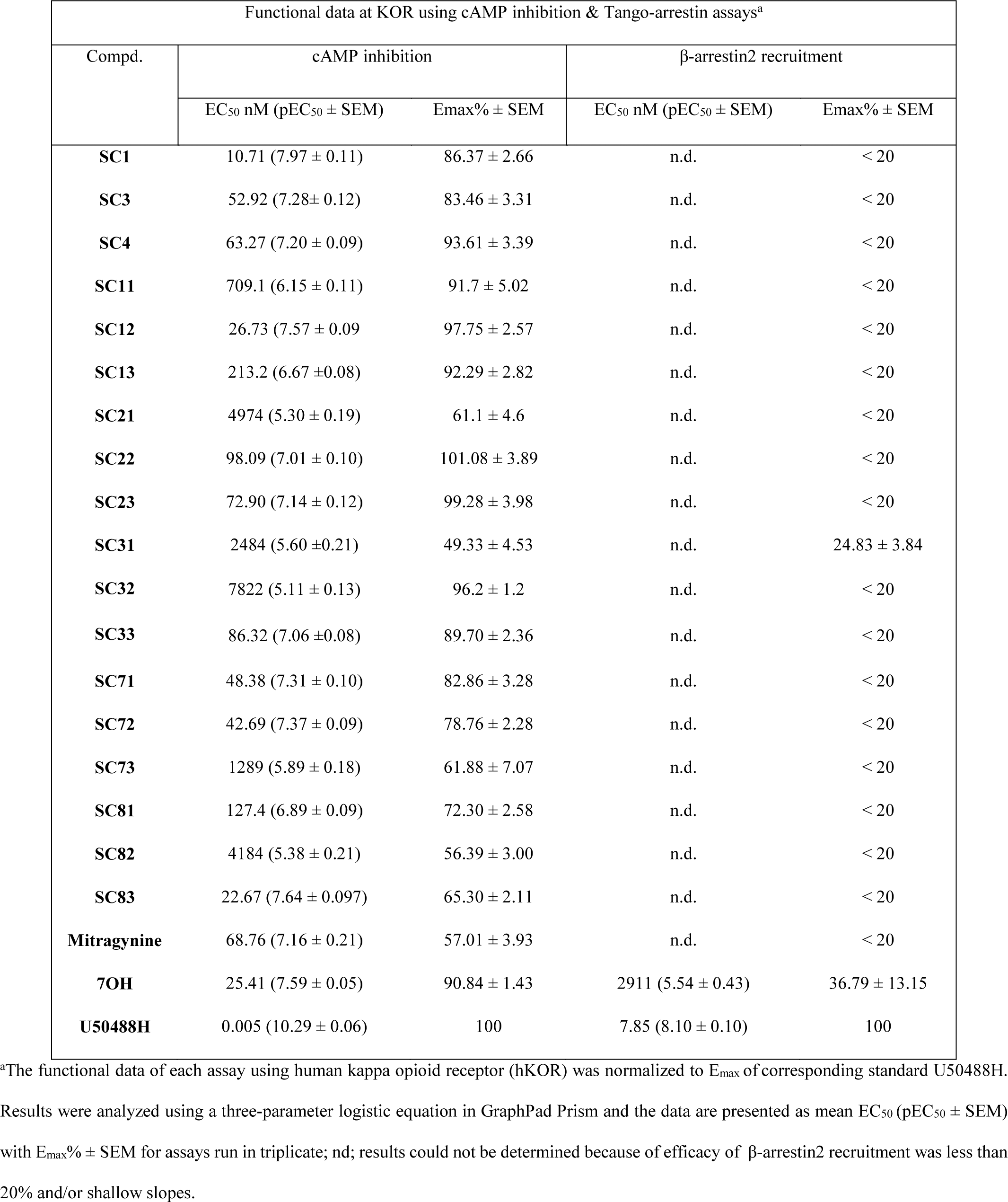
Functional studies at KOR using cAMP inhibition & Tango-arrestin assays.

**Appendix 1-Table 2.**
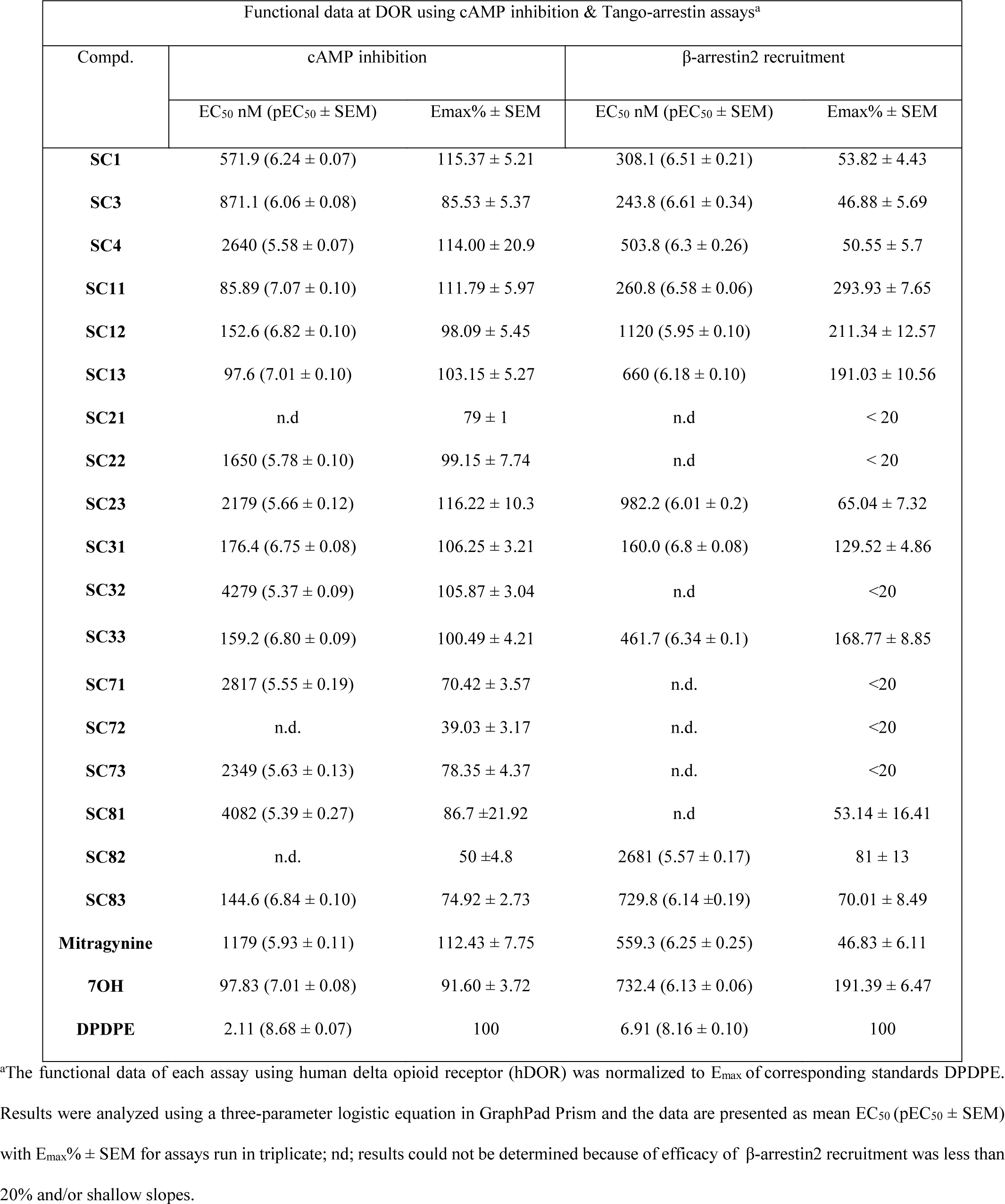
Functional studies at DOR using cAMP inhibition & Tango-arrestin assays.

**Appendix 1-Table 3.**
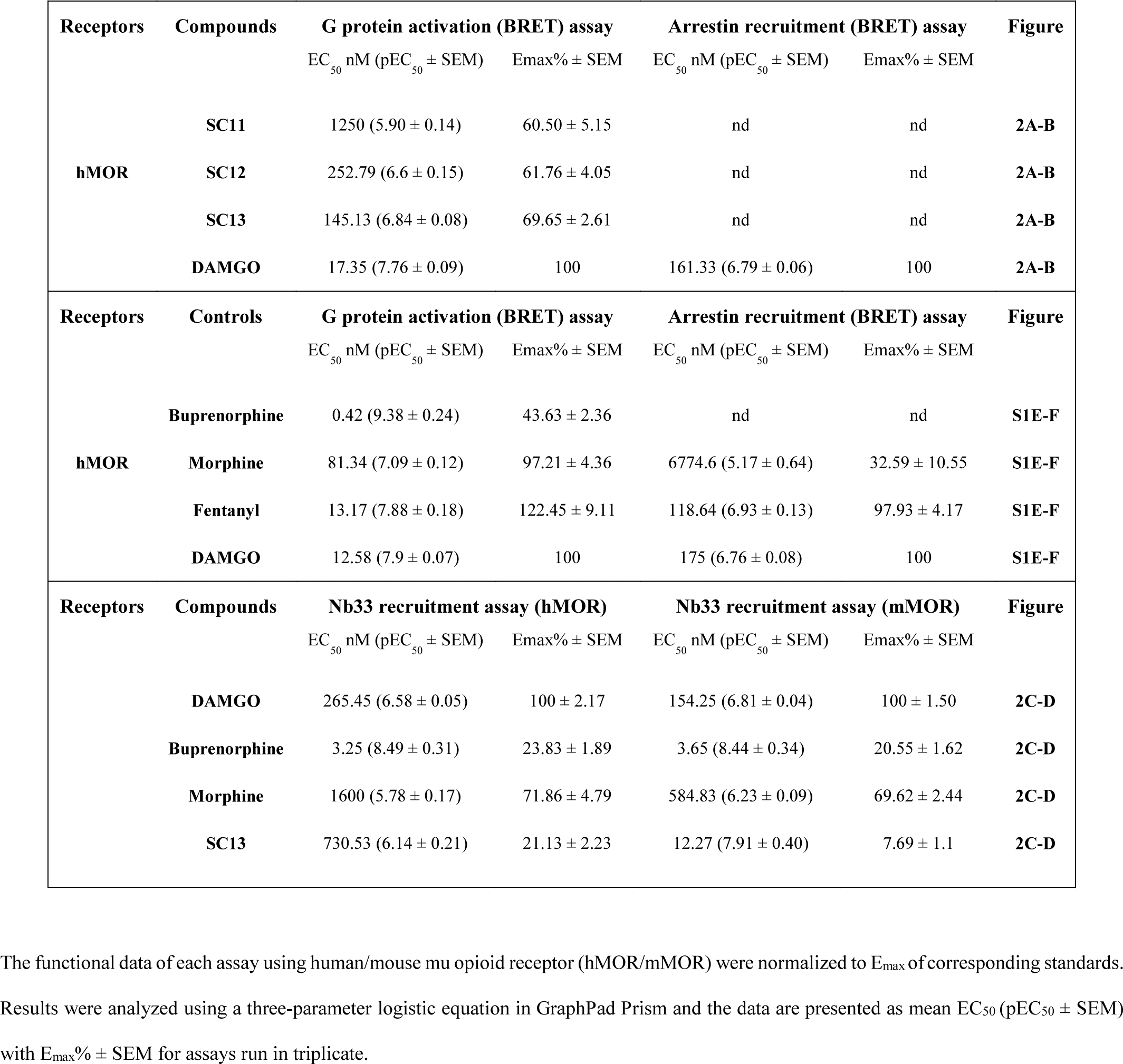
Functional binding data of the compounds at opioid receptors (hMOR/mMOR).

**Appendix 1-Table 4.**
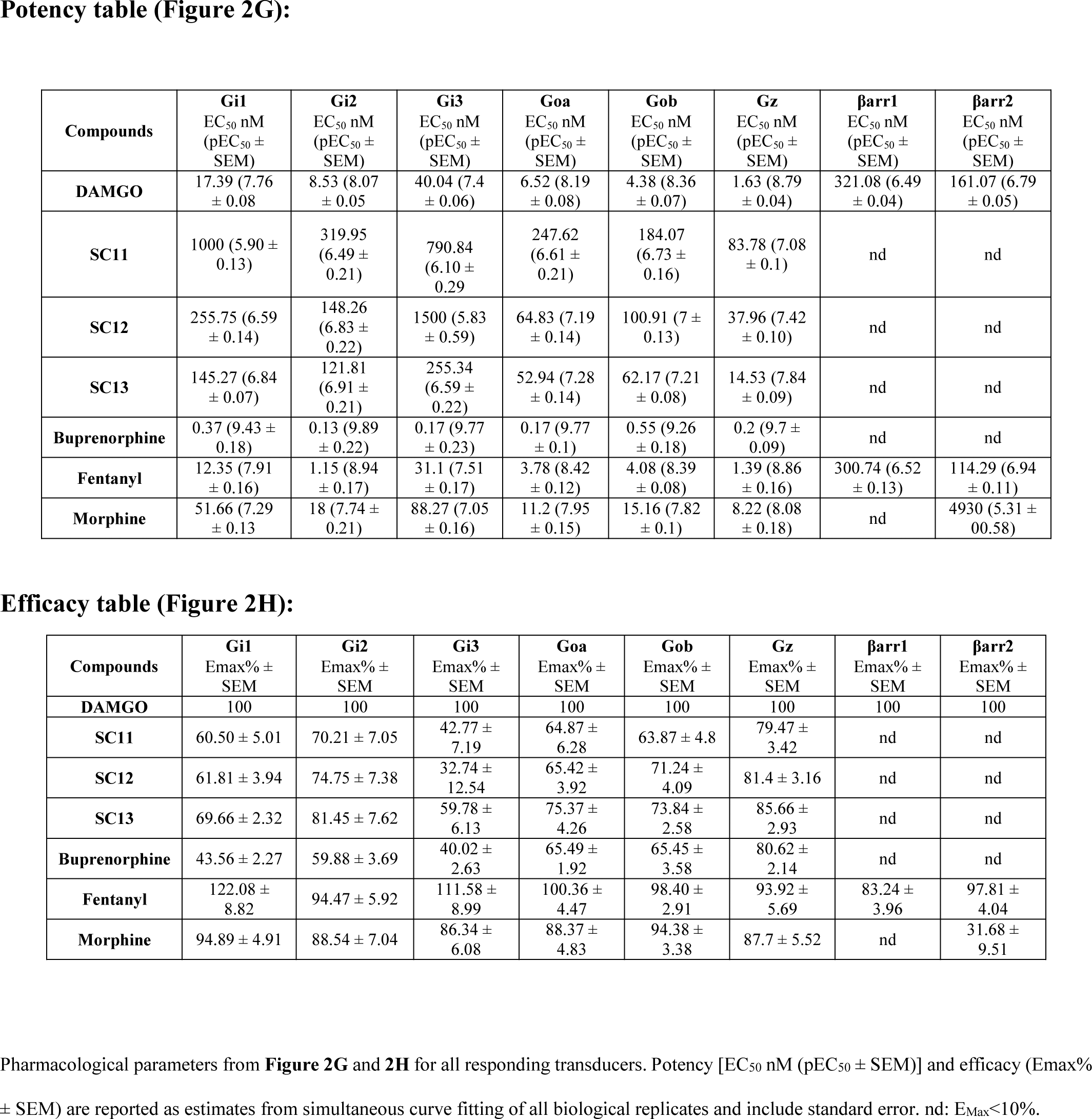
Potency and efficacy table for TRUPATH assay

**Appendix 1-Table 5.**
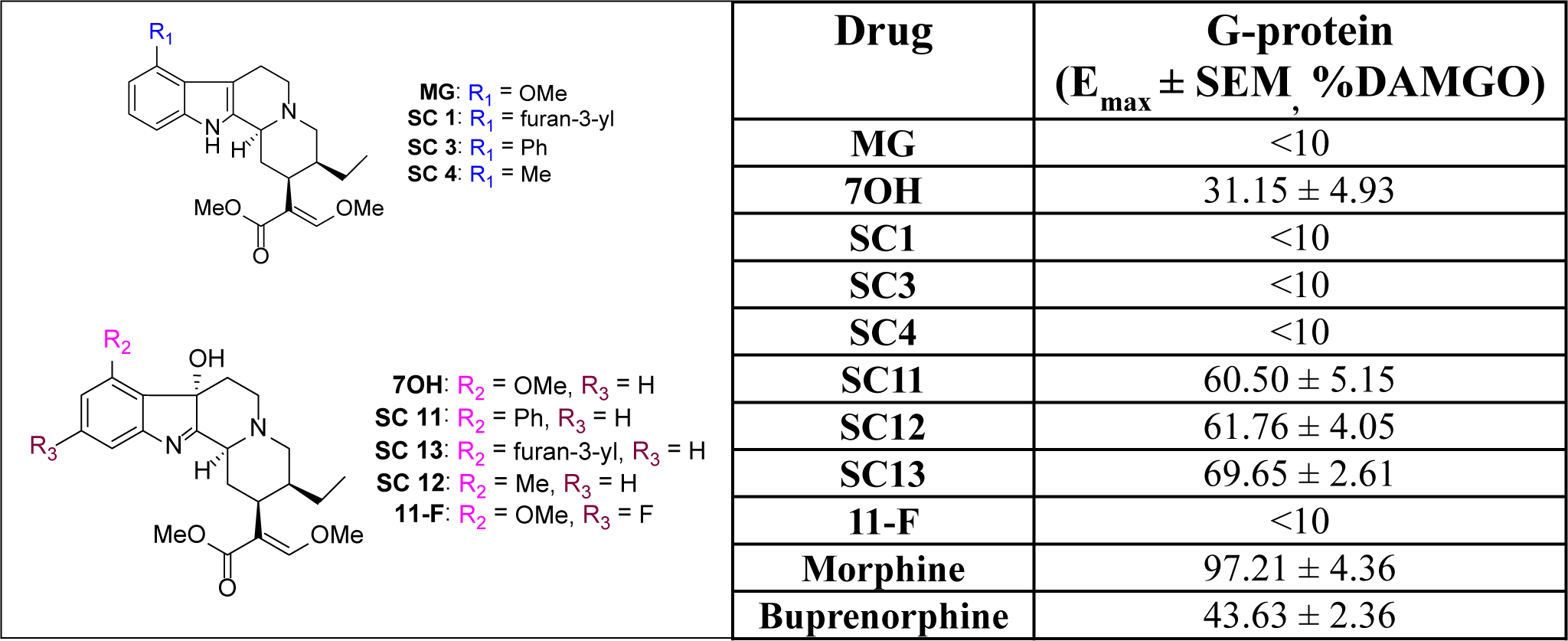
Compounds and data used to build the statistical models.

**Appendix 1-Table 6.**
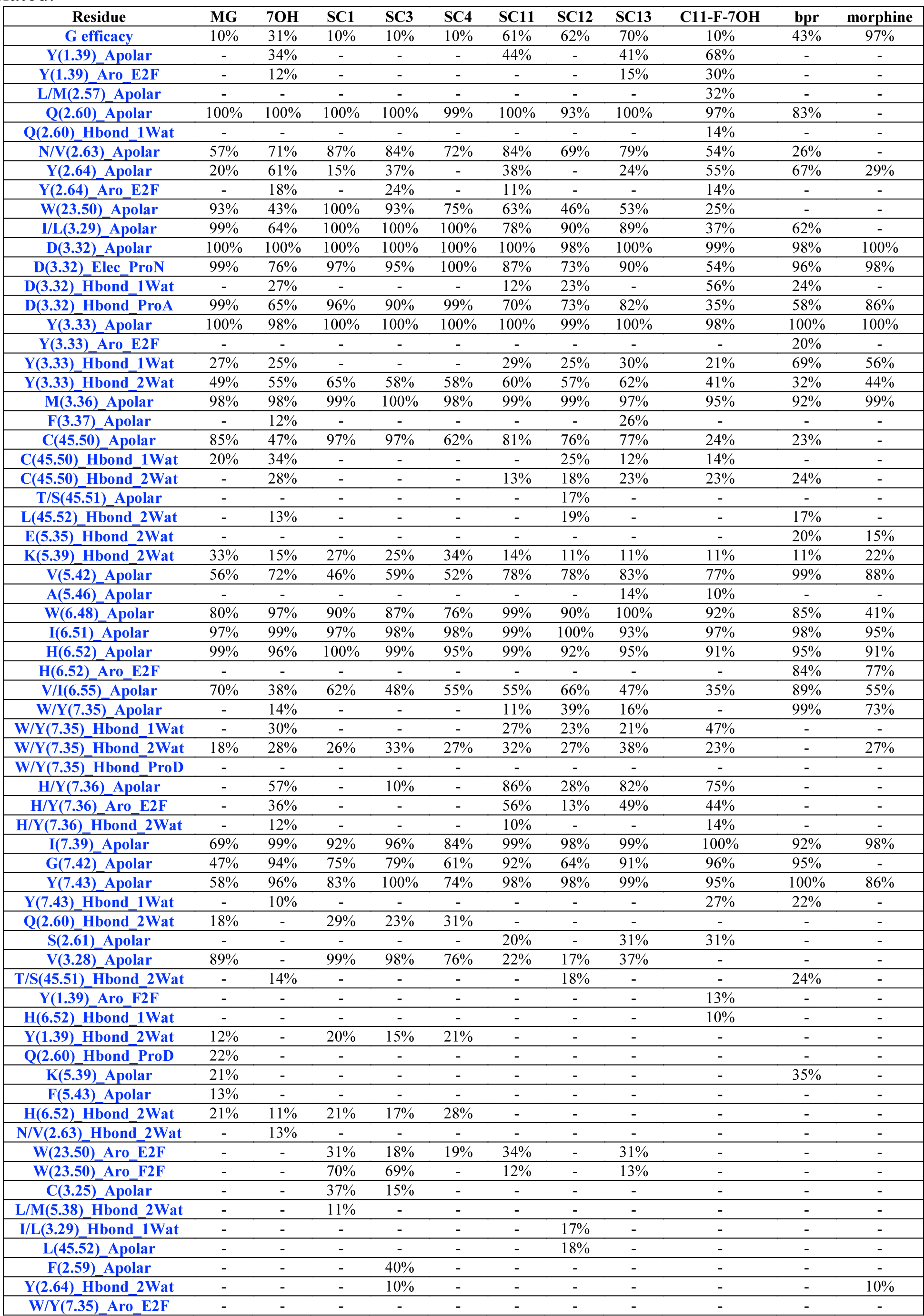
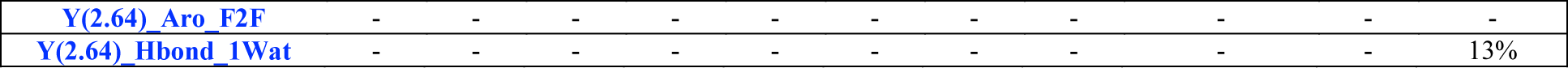
Average Structural Interaction Fingerprints (SIFt) probability for each ligand simulated.

**Appendix 1-Table 7.**
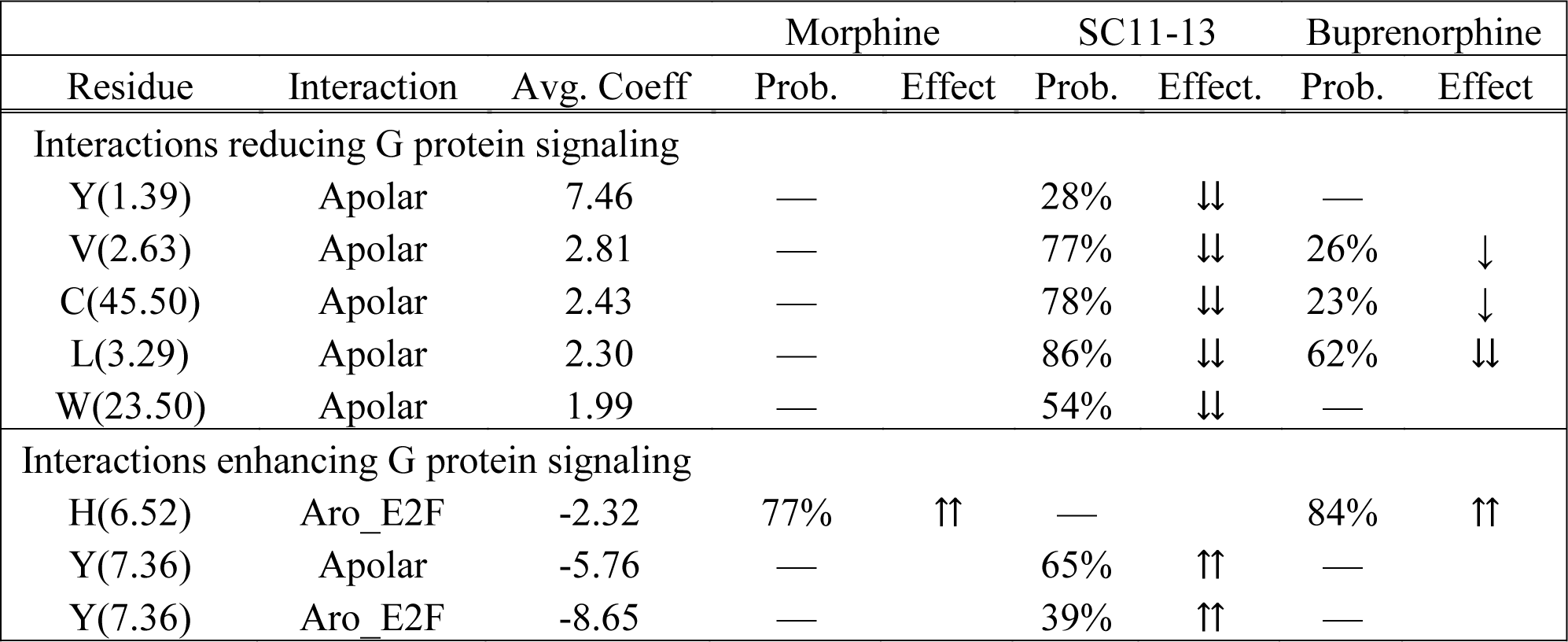
Interactions in the top statistical models that are predicted to either enhance (negative coefficients) or reduce (positive coefficients) ligand-induced MOR activation and consequent G protein signaling.

**Appendix 1-Table 8.**
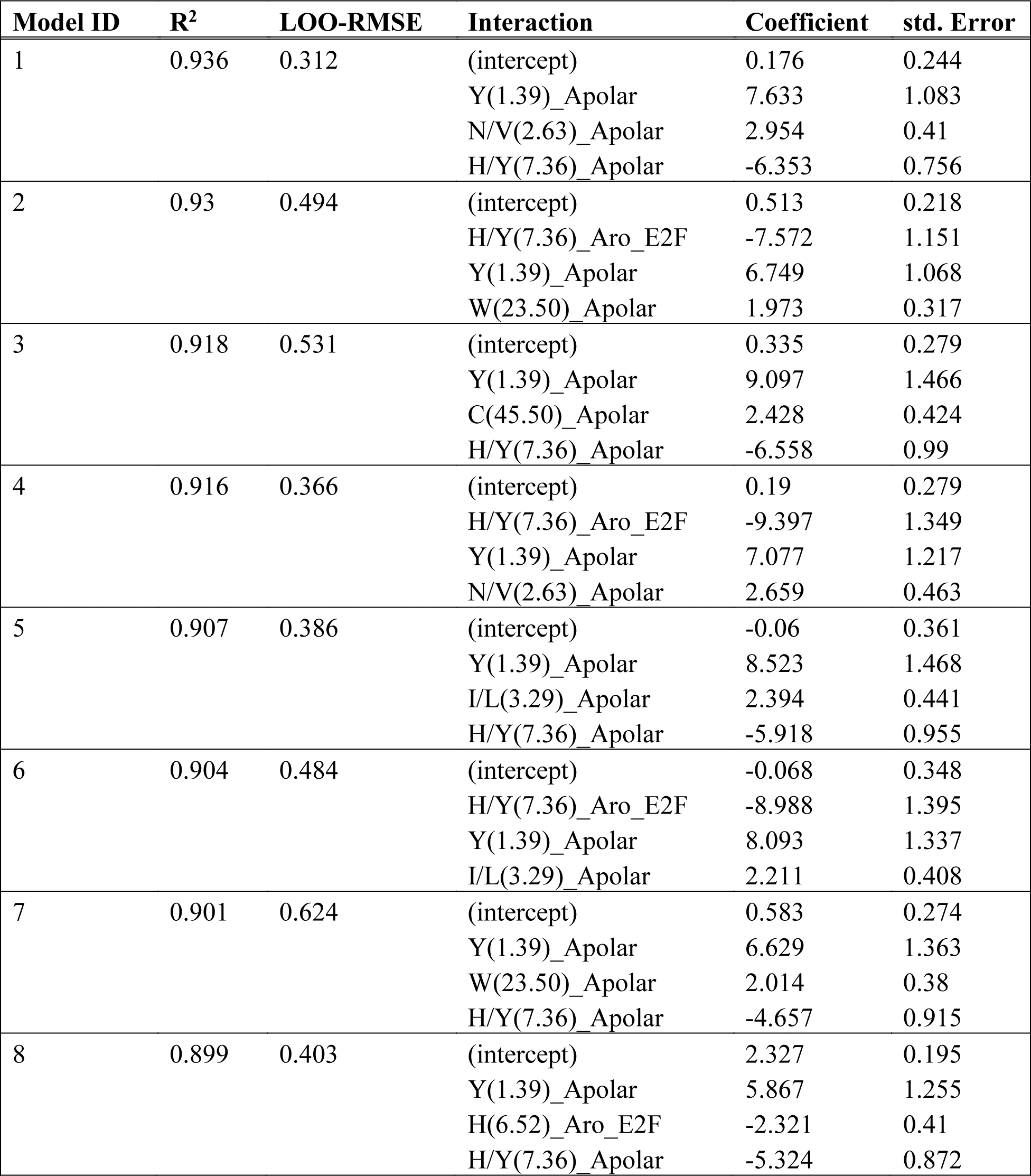
Selected models for the prediction of the negative log of the efficacy -log(E_Max_) as a function of interaction probabilities. The R^2^ on the full training set and the LOO-RMSE are reported for each model, as well as the values of the coefficient estimates and their standard errors.

**Appendix 1-Figure 1.**
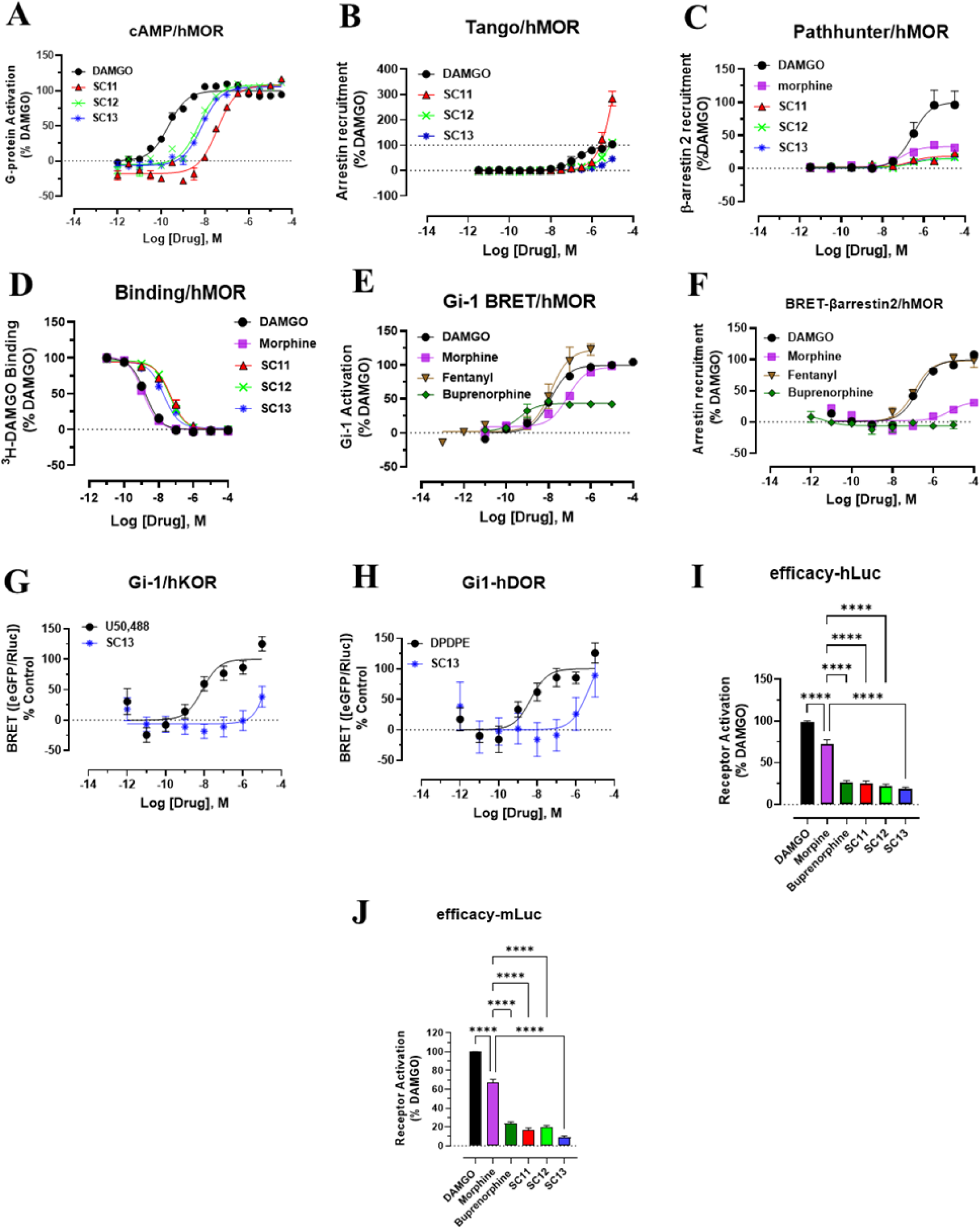
Characterization of SC11-13 in cAMP, Tango arrestin, PathHunter arrestin, binding, Nb33 recruitment at h/mMOR and Gi-signaling at hKOR and hDOR. **A) SC11**-**13** are full agonists at hMOR in cAMP inhibition (N=3) compared to DAMGO. See table **1** (main paper) for values **B) SC11-13** showed robust arrestin recruitment (E_max_>100% for **SC11-12** &45% for **SC13**) with poor potency (EC_50_>10µM) in Tango assays at MOR. See table **1** (main paper) for values. **C) SC11**-**13** in PathHunter- arrestin recruitment assays (n=3) show less βarrestin-2 recruitment compared to DAMGO. **β-arrestin2: SC11** EC_50_ nM (pEC_50_ ± SEM) = n.d., E_max_% ± SEM = <20%, **β-arrestin2: 2 SC12** EC_50_ nM (pEC_50_ ± SEM) = n.d., E_max_% ± SEM = <20%, **β-arrestin2: SC13** EC_50_ nM (pEC_50_ ± SEM) = n.d., E_max_% ± SEM = <20%, **β-arrestin2: morphine** EC_50_ nM (pEC_50_ ± SEM) = 80.08 (7.09 ± 0.17) nM, E_max_% ± SEM = 33.08 ± 1.84, **β-arrestin2: DAMGO** EC_50_ nM (pEC_50_ ± SEM) = 281.57 (6.55 ± 0.2) nM. . **D)** In competitive radioligand binding assays in MOR-CHO using _3_H- DAMGO as radioligand, **SC11-13** labelled MOR with high to reasonable affinity. **SC13** K_i_ (pK_i_ ± SEM) = 6.05 (8.22± 0.08), **SC12** K_i_ (pK_i_ ± SEM) = 12.33 (7.91 ± 0.03), **SC11** K_i_ (pK_i_ ± SEM) = 15.42 (7.81 ± 0.06), **morphine** K_i_ (pK_i_ ± SEM) = 0.37 (9.42 ± 0.04), **DAMGO** K_i_ (pK_i_ ± SEM) = 0.49 (9.31 ± 0.03). **E)** Gi-1 activation in BRET assays of controls. Fentanyl had higher efficacy over DAMGO. Efficacy of morphine was 94% and buprenorphine showed 44% efficacy. See **appendix 1-table 3** for values. **F)** β-arrestin2 recruitment in BRET assays of controls. Fentanyl showed robust arrestin recruitment with efficacy of 94%, morphine showed 31% efficacy while buprenorphine showed no recruitment. See **appendix 1-table 3** for values. **G)** No measurable Gi-1 potency was observed for **SC13** at hKOR. **U50488** EC_50_ nM (pEC_50_ ± SEM) = 8.07 (8.09 ± 0.27) nM. **SC13** EC_50_ nM (pEC_50_ ± SEM) = n.d., E_max_% ± SEM = 38 ± 17. **H)** No measurable Gi-1 potency was observed for **SC13** at hDOR. **SC 13** EC_50_ nM (pEC_50_ ± SEM) = n.d; E_max_% ± SEM = 88 ± 35 at hDOR. **I)** Efficacy of **SC** compounds, buprenorphine and morphine at the human opioid receptors in BRET-based Nb33 recruitment assaysare shown as a percentage of receptor activation relative to the full agonist, **DAMGO**. **SC11-13** had significantly lower efficacy than DAMGO (p<0.0001) and morphine (p<0.0001) and similar efficacy to buprenorphine. Statistical significance was determined using one-way ANOVA followed by Dunnett’s multiple comparison test, F(5,68)=239.172.5, p<0.0001. **J)** Efficacy of **SC** compounds, buprenorphine and morphine at the mouse opioid receptors in BRET-based Nb33 recruitment assays are shown as a percentage of receptor activation relative to the full agonist, **DAMGO**. **SC11-13** had significantly lower efficacy than DAMGO (p<0.0001), morphine (p<0.0001), and similar efficacy to buprenorphine. Statistical significance was determined using one-way ANOVA followed by Dunnett’s multiple comparison test, F(5,64)=572.5, p<0.0001.

**Appendix 1-Figure 2.**
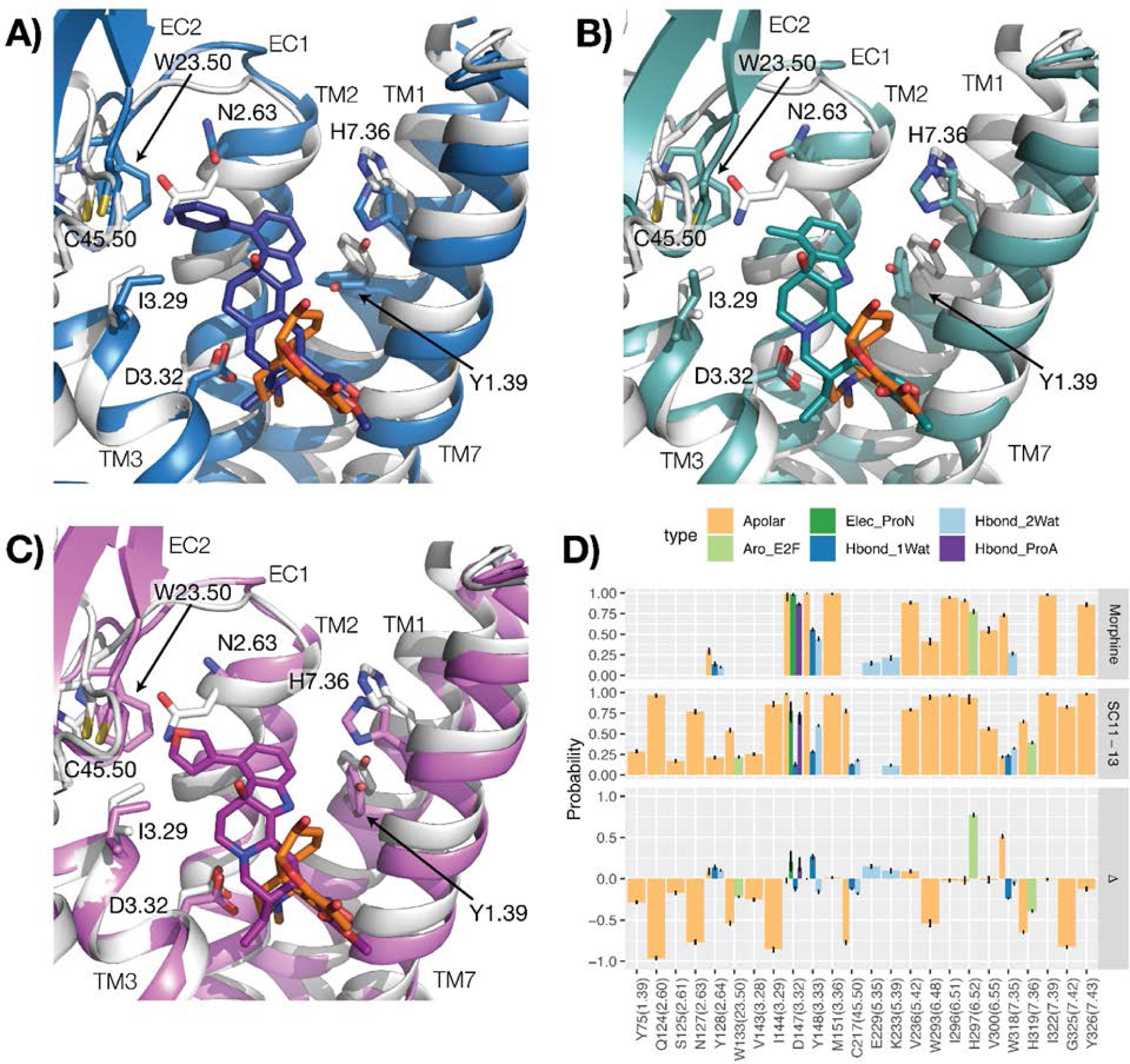
Binding modes and interactions of SC11-13 compared to morphine. (A-C) Representative conformations of the most populated clusters from MD simulations of MOR bound to SC11 (blue), SC12 (teal), and SC13 (purple) (panels A-C, respectively), compared to a representative conformation of MOR bound to morphine (orange). The protein is represented as a gray cartoon in the morphine-MOR complex. Residues identified in the best 8 performing models on experimental data are indicated with sticks. Transmembrane helices 5 and 6 are not shown for clarity. (D) Differences (plot at the bottom) between average structural interaction fingerprints (SIFts) calculated for SC11-13 (plot in the middle) and SIFts calculated for morphine (plot at the top).

**Appendix 1-Figure 3.**
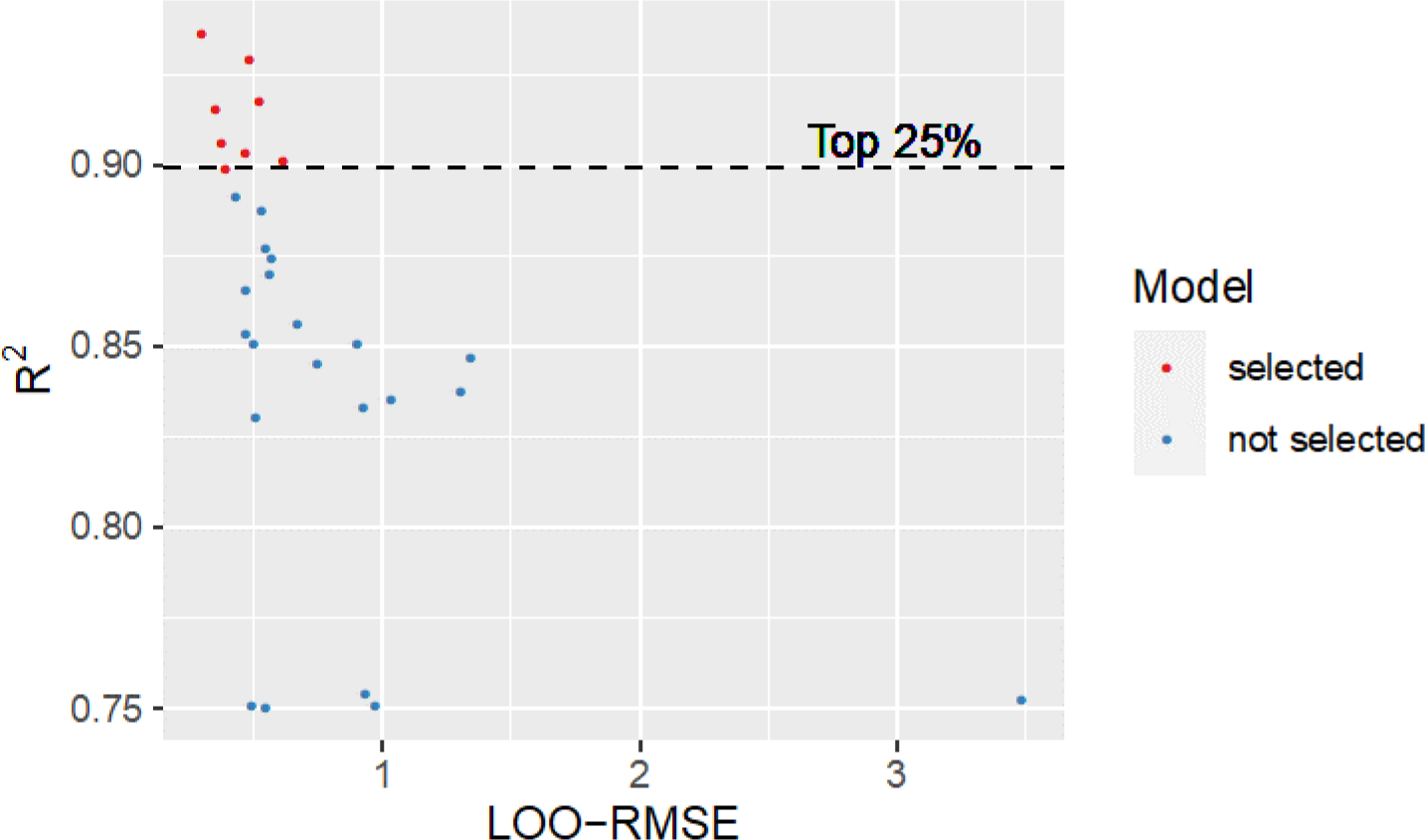
Full training set R_2_ validation and leave-one-out (LOO) cross-validation root mean square error (RMSE) for models with R^2^>0.75. Models with R^2^ in the top quartile (red points) were selected as best performing models on experimental data.

**Appendix 1-Figure 4.**
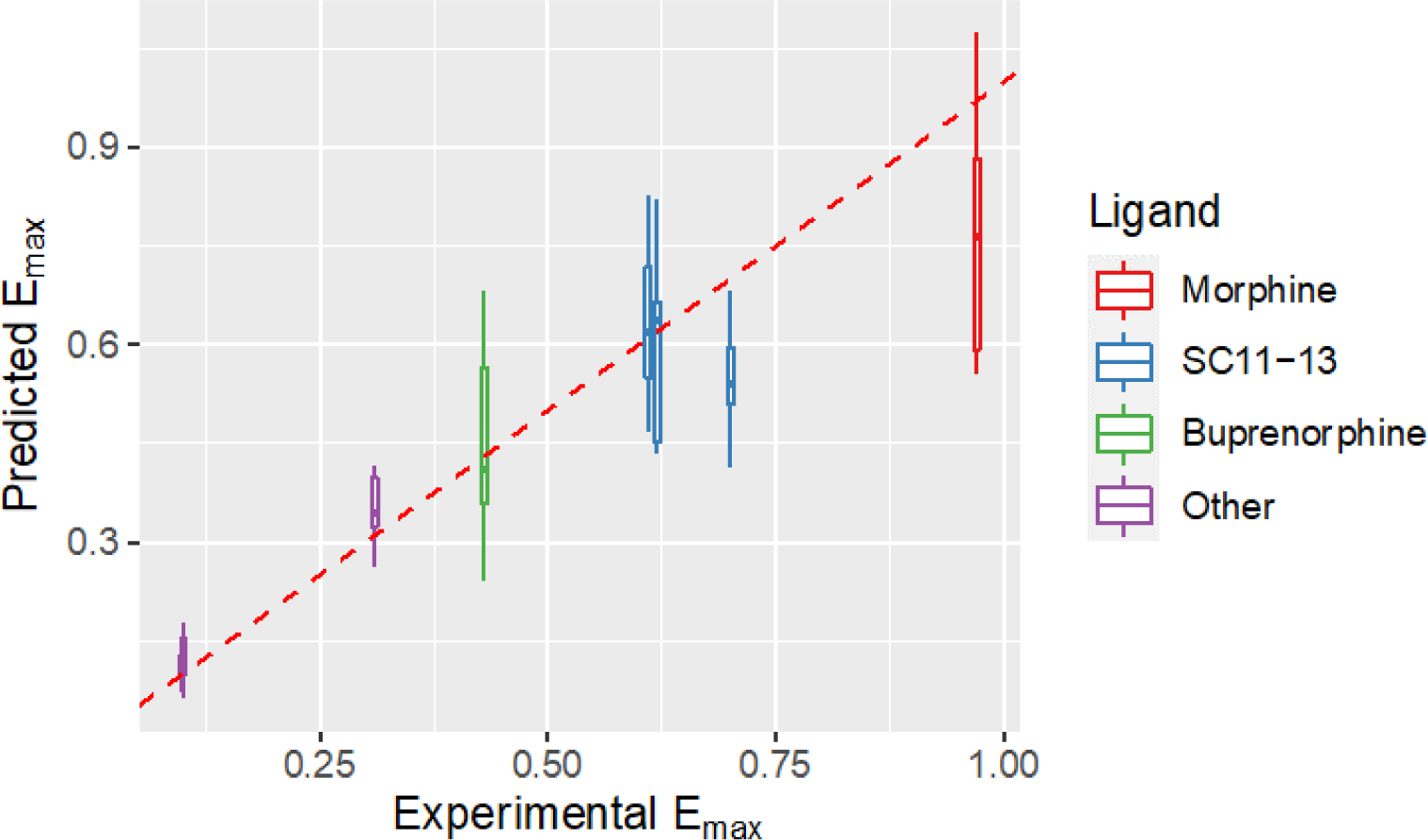
Values of the negative logarithm of the G protein efficacy E_max_ predicted from the selected top 25% models, compared to the experimental values for **morphine** (red), the **SC11-13** ligands (blue), **buprenorphine** (green), and the remaining 6 ligands in the training set (purple).

